# Cellular determinants of parvovirus B19 infection in the human placenta

**DOI:** 10.1101/2025.09.12.675789

**Authors:** Corinne Suter, Melanie Küffer, Jan Bieri, Amal Fahmi, David Baud, Marco P. Alves, Carlos Ros

## Abstract

Parvovirus B19 (B19V) is a prevalent human pathogen that can cross the placenta by a mechanism that remains unknown, posing a risk of severe fetal complications, particularly during the first trimester of pregnancy. We investigated the expression of B19V-specific receptors in the three trophoblast cell types, cytotrophoblasts (CTBs), syncytiotrophoblasts (STBs), and extravillous trophoblasts (EVTs), and assessed their susceptibility to infection. VP1uR, the erythroid-specific receptor that mediates viral uptake and infection in erythroid progenitor cells, is expressed in CTBs and STBs, but not in EVTs. Globoside, a glycosphingolipid that is essential for the escape of the virus from endosomes, is also expressed in these cells, except for choriocarcinoma-derived CTBs. In the latter, the absence of globoside can be overcome by promoting endosomal leakage with polyethyleneimine. While erythropoietin receptor (EpoR) signaling is associated with the strict erythroid tropism of B19V, it is not required for infection in trophoblasts. Transfection experiments revealed that highly proliferative first-trimester CTBs are more permissive to B19V infection than the low-proliferative CTBs from term placenta. These findings demonstrate that B19V targets and infects specific trophoblast cells, where viral entry and replication are collectively mediated by VP1uR, globoside, and high cellular proliferative activity, but are independent of EpoR signaling.

**Highlights:**

- Trophoblasts express the specific receptors necessary for B19V entry.
- Susceptibility and permissiveness to B19V vary by trophoblast subtype and gestational age.
- Highly proliferative first-trimester cytotrophoblasts show increased permissiveness to B19V.
- B19V infection in trophoblasts depends on globoside but is independent of EpoR signaling.

## Introduction

Parvovirus B19 (B19V) is a highly prevalent human pathogen responsible for *erythema infectiosum*, a pediatric disease marked by a characteristic erythematous rash and mild systemic symptoms^1^. The virus exhibits a pronounced tropism for erythroid progenitor cells (EPCs) within the bone marrow, where it establishes a lytic infection that disrupts normal erythropoiesis^2^. The lytic infection in EPCs leads to a transient interruption of erythropoiesis, which leads to the hematological disorders associated with the infection, particularly in individuals with chronic hemolytic anemia or immunosuppression. Infection during pregnancy is frequently associated with perinatal complications^3^. Despite its high prevalence and impact on human health, B19V remains a neglected viral infection, with no vaccines or antiviral treatments currently available.

The 25-nm capsid of B19V is assembled from only two proteins, VP1 and VP2^4^. The VP1 protein contains an N-terminal extension, termed the VP1 unique region (VP1u), which includes the receptor-binding domain (RBD)^5,6^. The VP1u cognate receptor, known as VP1uR, is a highly restricted receptor that has been exclusively found in the target EPCs in the bone marrow, explaining the marked erythroid cell tropism of the virus^7,8^. Transmission through the respiratory route is not mediated by VP1uR. Instead, the virus engages globoside, a glycosphingolipid expressed on the surface of the ciliated epithelial cells. The interaction with globoside is tightly regulated by pH^9^. Within the acidic environment of the nasal mucosa, the virus binds to globoside on the apical surface of epithelial cells. Subsequently, the virus is transported across the cell via transcytosis and released at the basolateral side, where the neutral pH facilitates its dissociation from globoside^10^. In addition to mediating viral translocation across the respiratory epithelium, globoside has an additional critical role. Following VP1uR-mediated endocytosis in EPCs, viral particles accumulate in endosomes, where low pH induces dissociation from VP1uR and binding to globoside, a critical interaction that facilitates escape from the endosomal compartment^9,11^.

The acute phase of infection is typically associated with exceptionally high-titer viremia, which is unparalleled by any other viral infection. During this highly viremic phase, the virus can invade the placenta and reach the fetus^12^. Transplacental transmission can result in severe complications, including fetal anemia, spontaneous miscarriage, *hydrops fetalis*, and intrauterine demise^13,14^. Early-to mid-pregnancy maternal infection markedly increases the chance of adverse effects on the fetus^15–17^. The increased risk in early gestation is attributed to the high expression of globoside in trophoblasts during this period^18^. Between 30-50% of women that become pregnant are nonimmune and susceptible to B19V infection^19–21^. In North America, between 1% and 2% of pregnant individuals become infected, with rates surging to around 10% during epidemic periods^21^. Among those with acute infection, transplacental transmission to the fetus has been documented in 17-51% of cases^22–24^. It has been estimated that approximately 3% of first-trimester spontaneous abortions may be due to B19V infection^25^. Pregnant women infected with B19V require careful monitoring, including serial ultrasounds to detect fetal anemia and *hydrops fetalis*. Early detection allows for timely interventions, such as intrauterine transfusions, to manage fetal anemia^21^.

Several studies have examined the interaction between B19V and the human placenta. Placental tissues from B19V infected women showed an increase in CD3-positive T cells and interleukin-2 production, suggesting that the infection triggers an immune response in the placenta^26^. The virus induces apoptosis in trophoblasts, which may contribute to placental dysfunction and adverse pregnancy outcomes^27^. Additionally, B19V has been linked to the deregulation of apoptotic pathways in placental tissues^28^. Empty capsids of B19V have been shown to bind to villous trophoblast cells via the globoside receptor, suggesting a role for globoside in facilitating viral transmission across the maternal-fetal interface^29^. A study reported a potential association between maternal B19V infection and placental abruption, suggesting that virus-induced apoptosis in trophoblasts may contribute to placental damage^30^. While these studies reveal the association between the virus and adverse pregnancy outcomes, none of them provide conclusive evidence of parvovirus B19 infection within the placenta, nor do they clarify the specific receptors involved or the cellular tropism of the virus among the different trophoblast populations.

Limited knowledge of B19V-host interactions at the maternal-fetal interface, combined with the lack of vaccines or antiviral therapies, places infected pregnant individuals at significant risk. Identifying the cellular determinants that regulate trophoblast permissiveness to B19V is crucial for advancing our understanding of the mechanisms driving vertical transmission. To address this, we employed trophoblast models representing the specialized trophoblast populations of the human placenta. Our findings reveal that VP1uR, the highly restricted B19V receptor known to mediate viral uptake and infection in EPCs, is also expressed in trophoblasts, identifying a previously unrecognized susceptibility factor at the maternal-fetal interface. In addition to VP1uR, globoside expression and increased cellular proliferation were revealed as major factors governing B19V infection in trophoblasts.

## Results

### Placental trophoblasts express the B19V receptor VP1uR

To detect the expression of VP1uR, we employed C-terminally and N-terminally truncated VP1u constructs. The C-terminally truncated construct (ΔC128) retains the functional receptor-binding domain (RBD; aa 5 to 80), while the N-terminally truncated construct (ΔN29) carries a disrupted, non-functional RBD and served as a negative control (Fig. 1A). These FLAG-tagged VP1u constructs were incubated with UT7/Epo, a human megakaryoblastoid cell line that expresses VP1uR^6^; BeWo and JEG-3 cells, first-trimester choriocarcinoma-derived cytotrophoblasts (CTBs); hPTC^CTB^, CTBs derived from term placenta^31^; and HTR-8/Svneo, a model of extravillous trophoblasts (EVTs)^32^. REH cells, a B-cell precursor leukemia line lacking VP1uR expression^7^, served as a negative control. After 30 minutes of incubation at 37°C, cells were fixed and stained with an anti-FLAG antibody to detect internalization. The VP1u construct containing the functional RBD (ΔC128) was detected in all trophoblast cell types except HTR-8/SVneo. The VP1u construct with the non-functional RBD (ΔN29) showed no binding (Fig. 1B). These findings reveal, for the first time, the expression of the B19V-specific receptor VP1uR beyond the erythroid lineage.

**Figure 1.**
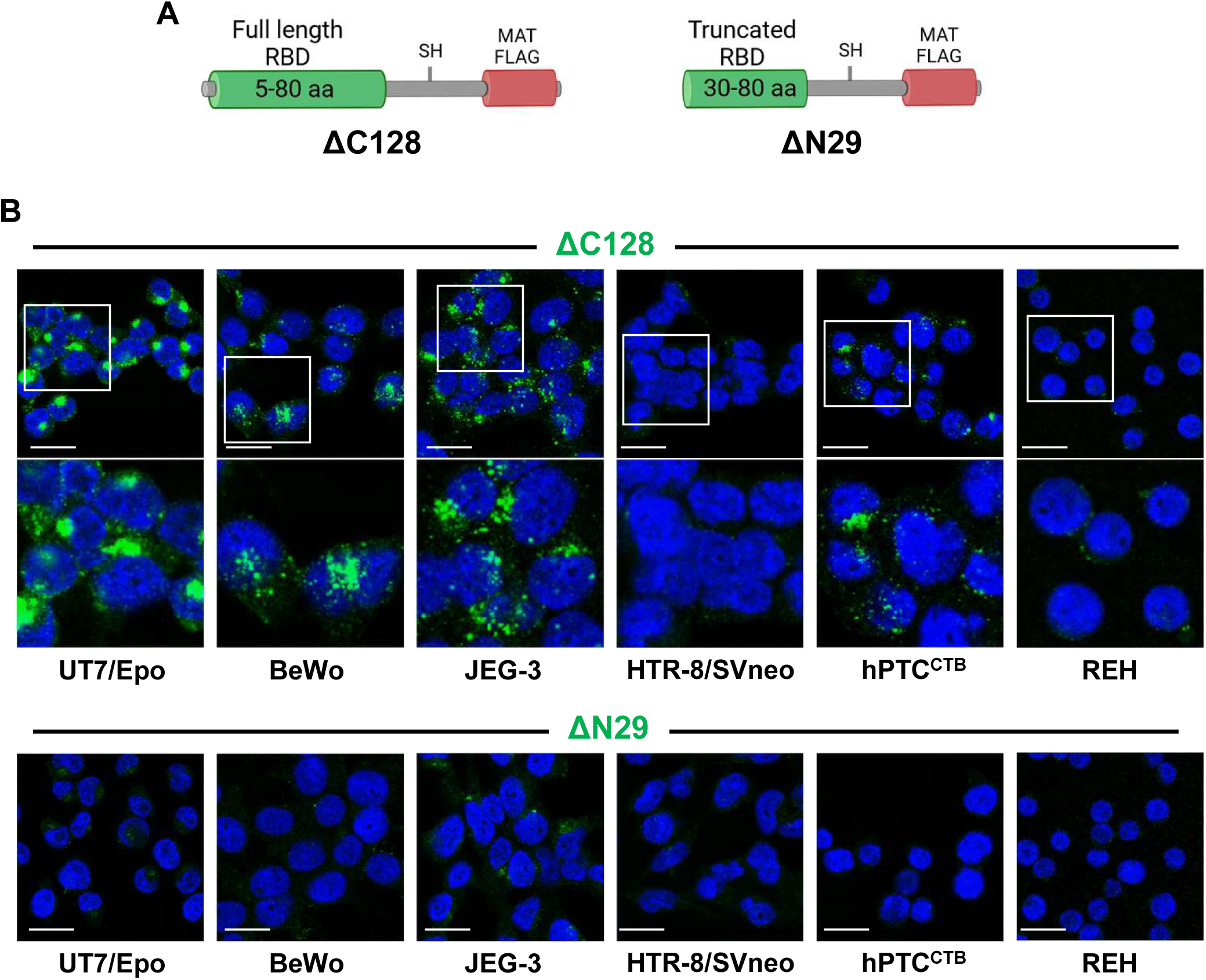
Detection of VP1uR expression in human placental trophoblasts. (A) Schematic representation of recombinant VP1u constructs. ΔC128 harbors a complete (5-80 aa) receptor-binding domain (RBD). ΔN29 harbors a truncated and non-functional RBD. The VP1u constructs include a cysteine-derived thiol group (-SH) for disulfide bond-mediated dimerization, an N-terminal MAT tag for efficient expression and solubility, and a C-terminal FLAG tag for detection with anti-FLAG antibodies. (B) Trophoblast cells were incubated at 37°C for 30 min with recombinant VP1u constructs and visualized by confocal microscopy with an anti-FLAG antibody (green). UT7/Epo cells served as positive and negative control, respectively. DAPI (blue). Scale bar, 20 μm.

Next, a virus-like particle (VLP)-based binding assay was used. The full-length B19V VP1u region was chemically conjugated to purified MS2 bacteriophage VLPs via click chemistry and labeled with Atto 488 for fluorescence detection. The use of VP1u-decorated MS2 VLPs allowed assessment of whether VP1u alone is sufficient to mediate the uptake of a virus-sized particle, independent of other B19V capsid domains or additional cellular factors. Since unmodified MS2 VLPs lack intrinsic affinity for trophoblasts, Atto 488-labeled MS2 VLPs without VP1u served as a negative control (Fig. 2A). MS2 VLPs constructs were incubated with the various trophoblast subtypes at 37°C for 30 minutes to allow receptor-mediated binding and internalization. UT7/Epo cells served as a positive control. Consistent with the recombinant VP1u binding assay, VP1u-coated MS2 VLPs bound to and were internalized by all trophoblast cell types tested, with the exception of HTR-8/SVneo. No binding was detected with Atto 488-labeled MS2 VLPs lacking VP1u (Fig. 2B). These results confirm the presence of functional VP1uR in CTBs from various placental origins and its absence in EVTs.

**Figure 2.**
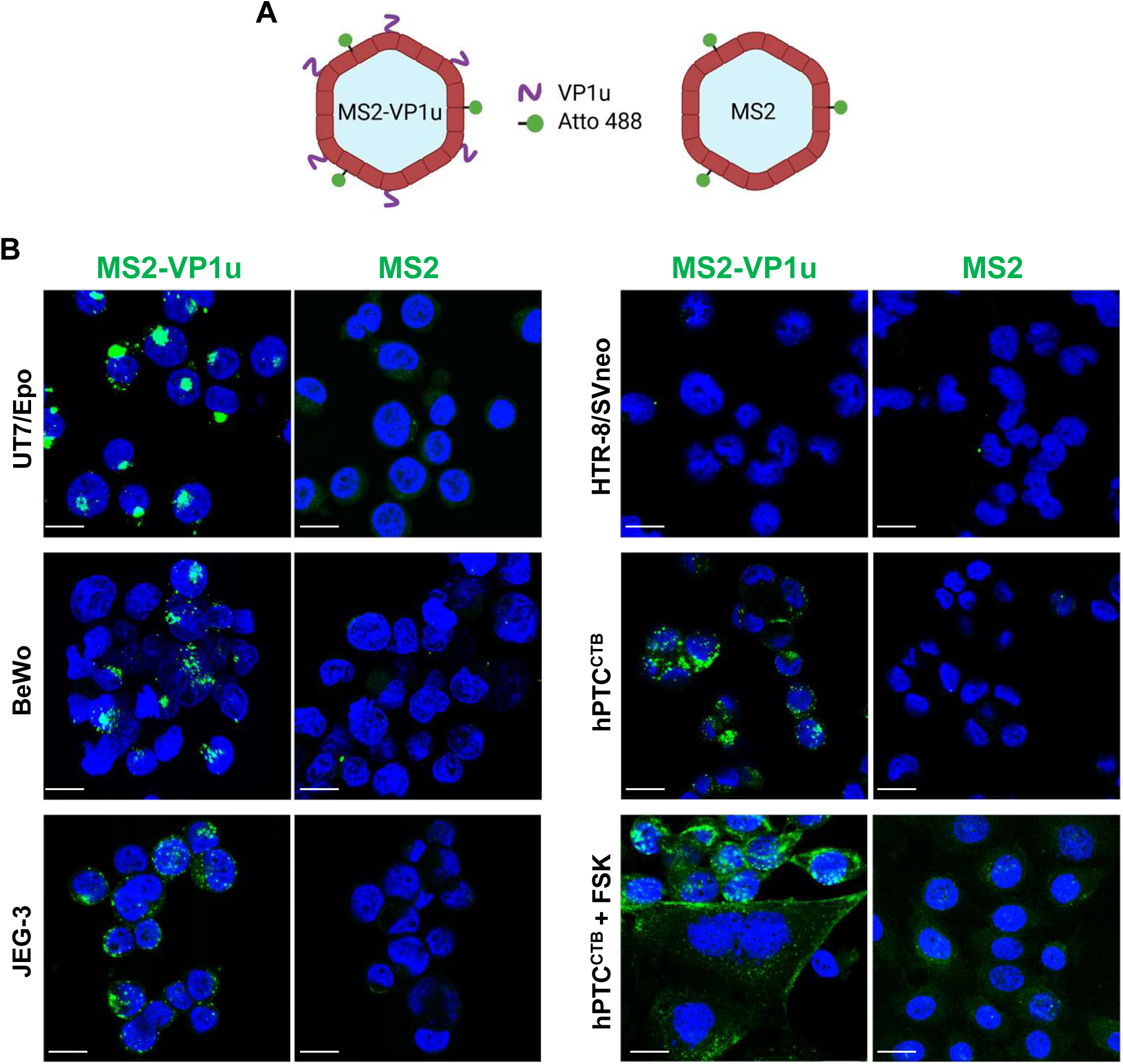
VP1uR is functional and mediates viral particle internalization in trophoblasts. (A) Schematic representation of MS2 VLPs conjugated with VP1u and/or Atto 488 by click chemistry. (B) UT7/Epo and different trophoblast subtypes were incubated at 37°C for 30 min with MS2 constructs and visualized by confocal microscopy. hPTC^CTB^ cells were treated with forskolin (hPTC^CTB^ + FSK) to induce differentiation into STBs. Following differentiation, cells were incubated at 37°C for 30 min with MS2 constructs and visualized by confocal microscopy. MS2-VLPs (green). DAPI (blue). Scale bar, 20 μm.

Forskolin induces the differentiation of CTBs into syncytiotrophoblasts (STBs) by activating adenylyl cyclase, which promotes cell fusion^33^. hPTC^CTB^ cells were differentiated into STBs using forskolin. STB differentiation was confirmed by the formation of large multinucleated syncytia. Fluorescence microscopy using MS2-VP1u VLPs showed that VP1uR expression was maintained following the differentiation of CTBs into STBs (Fig. 2B).

Given that the AXL receptor tyrosine kinase (AXL) has been proposed as a potential receptor mediating VP1u binding in erythroid cells^34^, we investigated whether its expression overlaps with VP1uR in placental trophoblasts. Our analysis revealed no consistent co-expression across trophoblast subtypes: BeWo and JEG-3 cells express VP1uR but not AXL; HTR-8/SVneo cells express AXL but lack VP1uR; and hPTC^CTB^ cells express both receptors (S1 Fig).

### Globoside expression in trophoblast cell lines

Globoside is essential for B19V infection, enabling transcytosis through the respiratory epithelium^10^, and facilitating endosomal escape of incoming virions in EPCs^11^. Given its essential role, we examined globoside expression across the different trophoblast subtypes to assess their potential susceptibility to B19V infection. The mRNA levels of globoside synthase (Gb4) and globotriaosylceramide synthase (Gb3), were determined by quantitative RT-PCR (RT-qPCR) using specific primers for β3GalNT1 (Gb4) and A4GalT (Gb3) mRNAs and normalized to GAPDH. UT7/Epo cells, which express globoside and are commonly used to study B19V infection, served as a positive control. Globoside knockout UT7/Epo cells, generated by CRISPR/Cas9-mediated knockout in a previous study^35^, and REH cells, which naturally lack globoside expression, were used as negative controls. The results showed that BeWo and JEG-3 cells expressed Gb3 synthase but lacked Gb4 synthase. In contrast, HTR-8/SVneo and hPTC^CTB^ cells expressed both Gb3 and Gb4 synthases (Fig. 3A). Immunofluorescence staining with a globoside-specific antibody confirmed the RT-qPCR results, showing that BeWo and JEG-3 cells lacked detectable globoside expression, whereas it was present in HTR-8/SVneo and hPTC^CTB^ cells (Fig. 3B).

**Figure 3.**
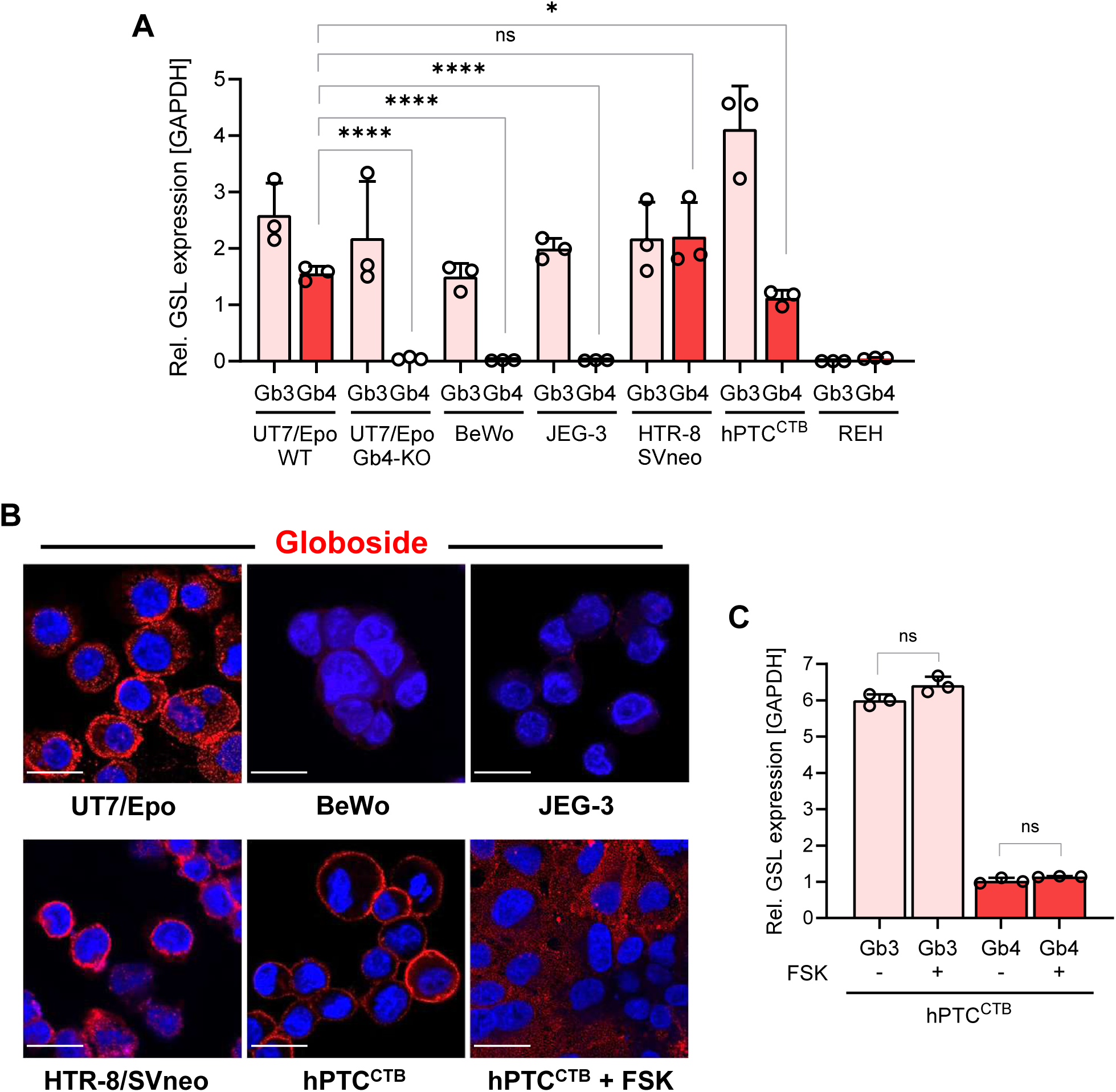
Globoside expression in different trophoblast subtypes. (A) Relative expression of Gb3 (Gb3 synthase) and Gb4 (Gb4 synthase) mRNA in different trophoblasts. mRNA levels were measured by RT-qPCR and normalized to GAPDH expression. GSL, glycosphingolipid. (B) Immunostaining of UT7/Epo and trophoblasts with an anti-globoside antibody (red). DAPI (Blue). Scale bar, 20 μm. (C) Relative expression of Gb3 and Gb4 mRNA in hPTC^CTB^ cells following differentiation into STBs using forskolin (FSK). mRNA levels were measured by RT-qPCR and normalized to GAPDH expression.

To examine whether globoside expression is maintained during the differentiation of CTBs into STBs, we induced syncytialization in hPTC^CTB^ cells using forskolin. Syncytialization induced with forskolin has been reported to alter the composition and abundance of the placental glycocalyx, potentially impacting glycosphingolipid expression^36^. Following STB differentiation, Gb4 expression remained detectable, indicating that globoside expression is maintained in STBs (Fig. 3B).

Consistently, the mRNA levels of Gb3 and Gb4 synthases remained unchanged before and after differentiation, indicating that syncytialization does not affect globoside expression (Fig. 3C). Furthermore, forskolin-induced syncytialization of BeWo cells did not restore globoside expression in these cells (S2 Fig. A). Considering that oxygen tension is low during early pregnancy and critically regulates gene expression^37^, and that Epo (erythropoietin) signaling through the Epo receptor (EpoR) in trophoblasts can influence differentiation and survival^38^, we investigated whether globoside expression is modulated by oxygen levels and/or Epo stimulation. BeWo cells were cultured under normoxic or hypoxic conditions, with or without Epo. While Gb3 expression remained consistent across all conditions, globoside expression was consistently undetectable, irrespective of oxygen tension or Epo stimulation (S2 Fig. B and C).

The expression of globoside was evaluated in cryosections of term placental tissue. To verify the presence of trophoblast subpopulations, sections were stained with BCL-2, Trop-2, and CD138. BCL-2 is used to label syncytiotrophoblast, Trop-2 identifies both cytotrophoblasts and syncytiotrophoblasts, and syndecan-1 marks the syncytiotrophoblast apical surface^39,40^. These markers together allow reliable delineation of trophoblast compartments in placental tissue sections. Staining with an antibody against globoside revealed a scattered signal distributed along the periphery of the villi and, more prominently, within the villous core. This pattern suggests that globoside expression is not restricted to the trophoblast layers but is more abundant in non-trophoblast cell populations residing within the villous core (S3 Fig). These findings corroborate previous studies that have reported widespread expression of globoside in placental trophoblasts^18,41,42^. The absence of globoside expression in the choriocarcinoma cell lines BeWo and JEG-3 is likely attributable to their malignant nature, as malignant transformation is known to alter glycosyltransferase expression^43,44^.

### B19V uptake by placental trophoblasts correlates with VP1uR expression

To assess the functional relevance of VP1uR expression in trophoblast cells, a viral internalization assay was performed. UT7/Epo cells were included as a positive control, and REH cells as a negative control to establish background signal. After 1 h incubation with B19V at 37°C, residual surface-bound virus was removed by trypsinization, and intracellular viral genomes were quantified by qPCR. B19V DNA was readily detected in BeWo, JEG-3, and hPTC^CTB^ cells, all of which express VP1uR, whereas no internalization above background was observed in VP1uR-negative HTR-8/SVneo and REH cells (Fig. 4A).

**Figure 4.**
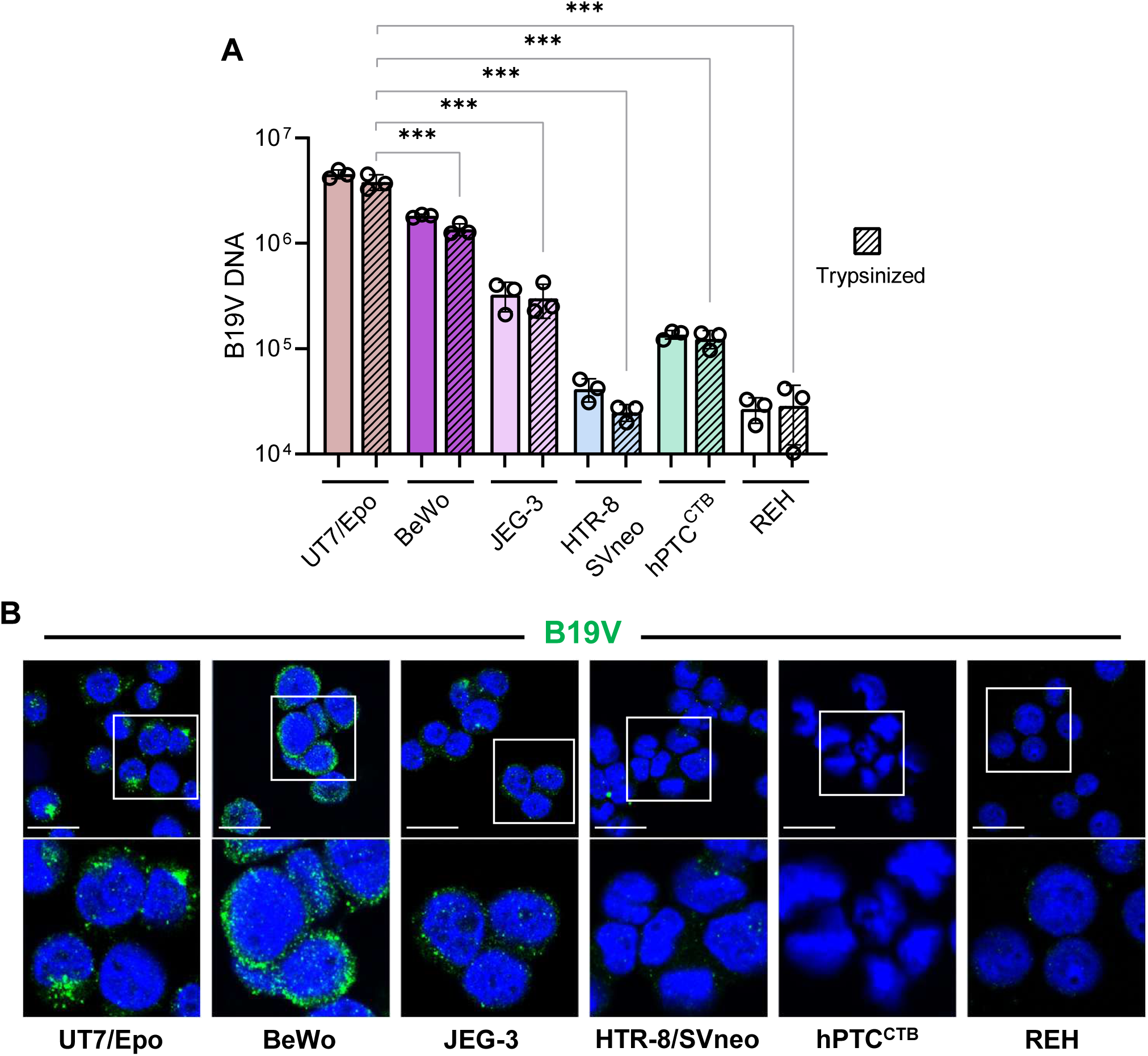
B19V uptake in trophoblasts correlates with VP1uR expression. (A) Cells were incubated with B19V (10^5^ genome equivalents per cell) at 37°C for 1 h. After incubation, some cells were washed to remove unbound viruses, while others were washed and trypsinized to remove surface-bound but not internalized viruses. Viral genomes were quantified by qPCR. UT7/Epo cells served as a positive control, and REH cells as a negative control to define background signal. Two-sided Student’s t-test. The results are presented as the mean ± SD of three independent experiments. ∗∗∗*p* < 0.001. (B) Immunofluorescence images showing B19V uptake in UT7/Epo and trophoblast cells. Cells were incubated with B19V (10^5^ genome equivalents per cell) at 37°C for 30 min. After incubation, cells were washed, fixed and stained with a monoclonal antibody against capsids (mAb 521, green). DAPI (Blue). Scale bar, 20 μm.

To corroborate these findings, immunofluorescence staining for B19V capsid proteins was performed. A strong viral signal was observed in BeWo cells, weaker staining in JEG-3 and hPTC^CTB^, and no detectable signal in HTR-8/SVneo (Fig. 4B). The relative signal intensity mirrored the qPCR data, and notably, these differences in viral uptake were consistent with the levels of VP1uR expression observed in these cells (Fig. 1 and 2). Importantly, B19V uptake was detected exclusively in cells expressing VP1uR, highlighting its essential role in mediating viral entry in trophoblasts.

### B19V infection across trophoblast types and gestational stages

To determine whether B19V internalization in trophoblast cells results in productive infection, the expression of the early viral transcript NS1 was analyzed 24 h and 48 h post-infection. After extracting total RNA, samples were treated with DNase I to eliminate residual genomic DNA and subjected to RT-qPCR using primers specific for B19V NS1 mRNA. UT7/Epo cells served as positive control, while REH cells, which are non-permissive to B19V, were used as negative control to define the background signal. In BeWo and JEG-3, NS1 mRNA levels were in the order of three logs lower than in UT7/Epo cells, consistent with the lack of Gb4 expression in these cell lines, which is essential for endosomal escape^11^. Similarly, HTR-8/SVneo cells, which do not express VP1uR, showed minimal NS1 expression, supporting the conclusion that insufficient viral entry limits infection in these cells. In contrast, hPTC^CTB^ cells, which express VP1uR and Gb4, exhibited significantly higher NS1 expression, though still lower than levels observed in UT7/Epo cells (Fig. 5A).

**Figure 5.**
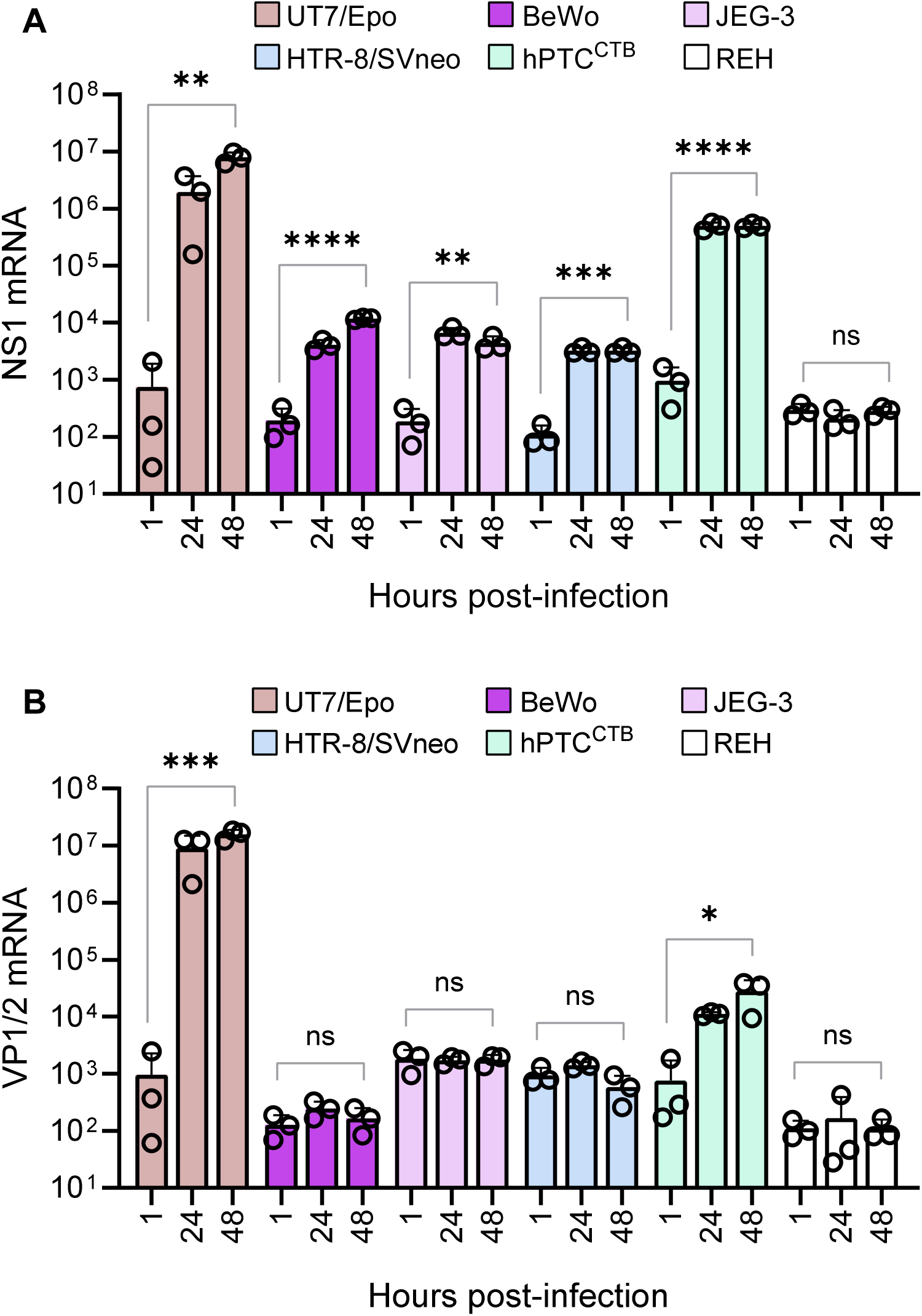
Expression of B19V non-structural and structural mRNAs in trophoblast cells. (A) Quantification of B19V infection in different trophoblast subtypes. UT7/Epo and REH cells were used as positive and negative controls, respectively. Cells were infected with B19V (10^5^ genome equivalents per cell) at 37°C for 30 minutes. NS1 mRNA levels were quantified by RT-PCR at 1 h (input), 24 h and 48 h post-infection. (B) VP1/VP2 mRNA levels were quantified in the same experimental setup to assess structural viral gene expression. All results are presented as the mean ± SD of three independent experiments. Statistical significance was calculated using two-sided Student’s t-test. ∗*p* < 0.05; ∗∗*p* < 0.01; ∗∗∗*p* < 0.001; ∗∗∗∗*p* < 0.0001; ns, non-significant.

Progression of infection beyond the early phase was evaluated by quantifying mRNA levels of the structural viral proteins VP1 and VP2 using RT-qPCR. Consistent with low or absent expression of NS1, VP1/2 transcripts were undetectable in BeWo, JEG-3, and HTR-8/SVneo cells, with values comparable to the input control. In contrast, VP1/2 mRNA was detected in hPTC^CTB^ cells, indicating that infection had progressed beyond the early transcriptional stage (Fig. 5B). However, VP1/2 transcript levels in hPTC^CTB^ remained substantially lower than those observed in UT7/Epo cells, suggesting limited efficiency of full viral replication.

B19V infection requires both VP1uR and Gb4 for cellular entry and depends on active cell proliferation for successful replication. Despite a robust NS1 expression, the limited viral structural mRNA expression observed in hPTC^CTB^ may be attributed, at least in part, to the reduced proliferative capacity and increased cellular senescence of CTBs during late gestation^45^. To investigate this possibility, we assessed the proliferation rates of different trophoblast subtypes by quantifying β-actin gene expression over a four-day culture period. As expected, the results confirmed that first-trimester trophoblasts exhibit high proliferative activity, whereas hPTC^CTB^ derived from term placenta show the lowest proliferation rate (S4 Fig).

### Highly proliferating cytotrophoblasts from first-trimester placenta exhibit a high level of permissiveness to B19V infection

To assess the permissiveness of different trophoblast subtypes to B19V infection independently of viral entry, transfection assays were conducted using purified single-stranded viral DNA genomes. NS1 and VP1/2 mRNA levels were quantified by RT-qPCR two days post-transfection. Each cell type was assessed for transfection efficiency by introducing a GFP-expressing plasmid and quantifying the proportion of GFP-positive cells via fluorescence microscope. UT7/Epo cells were used as reference control. Transfection efficiency varied among the trophoblast cell lines: HTR-8/SVneo exhibited the highest efficiency (4.5-fold higher than UT7/Epo), followed by BeWo (3.5-fold). In contrast, JEG-3 and hPTC^CTB^ showed transfection rates comparable to those of UT7/Epo cells (Fig. 6A). Remarkably, NS1 mRNA quantification revealed that first-trimester CTBs, specifically JEG-3 and BeWo cells, exhibited significantly higher NS1 expression levels, showing 42-fold and 12-fold increases, respectively, compared to UT7/Epo cells (Fig. 6B).

**Figure 6.**
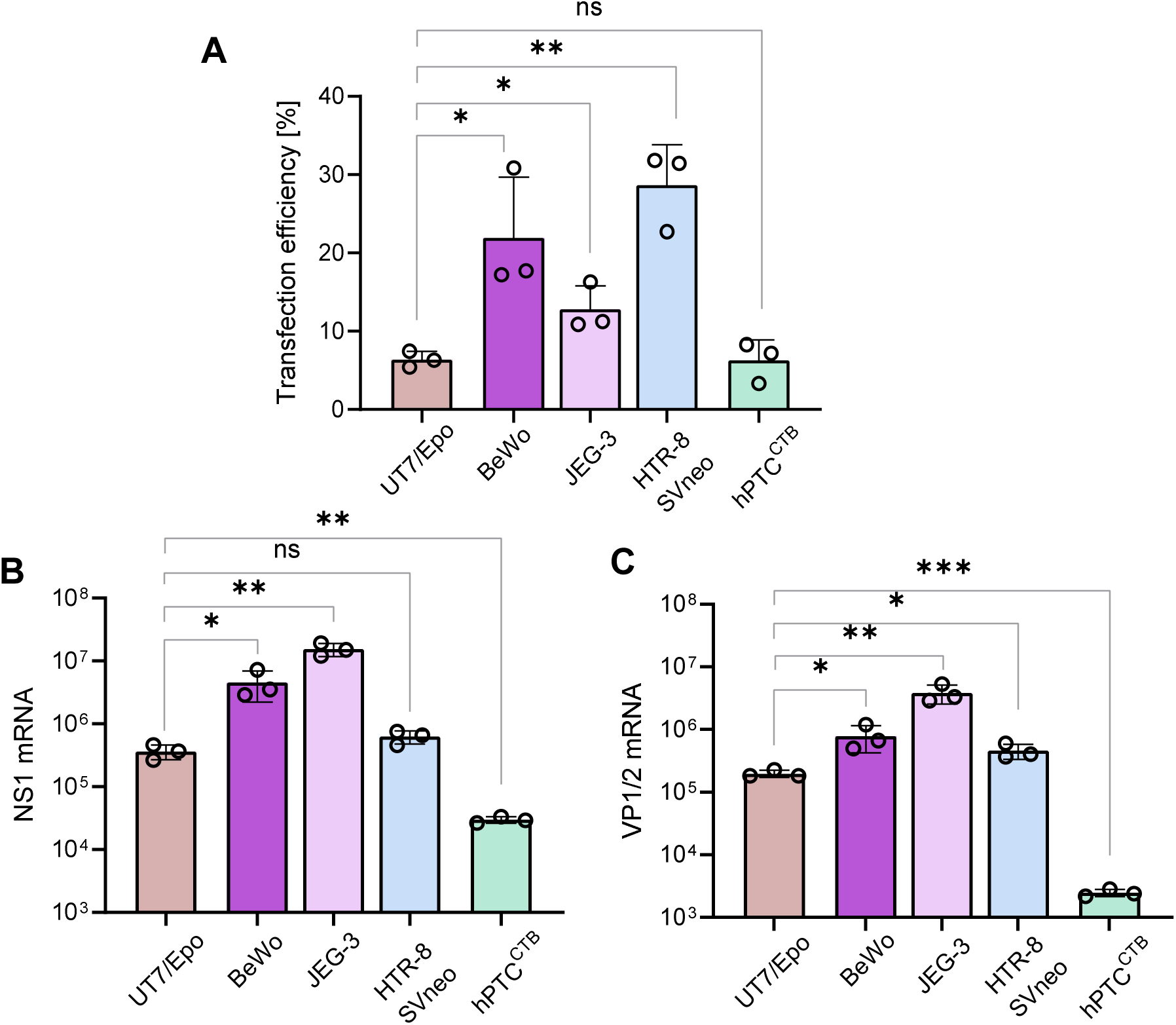
Permissiveness of trophoblast subtypes to B19V infection. (A) Transfection efficiency was evaluated with a plasmid encoding green fluorescent protein (GFP). Cells were visualized by fluorescence microscopy, and GFP-positive cells were quantified as a percentage of total cells. (B) Purified single-stranded B19V genomes were transfected into the indicated cell types. UT7/Epo cells were included as a permissive reference. At 24 h post-transfection, NS1 mRNA expression was quantified by RT-PCR to assess early viral gene expression. (C) Under the same experimental conditions as in (B), VP1/VP2 mRNA levels were quantified by RT-PCR to evaluate structural gene expression following transfection. All results are presented as the mean ± SD of three independent experiments. Statistical significance was calculated using two-sided Student’s t-test. ∗*p* < 0.05; ∗∗*p* < 0.01; ∗∗∗*p* < 0.001; ns, non-significant.

Given that mRNAs encoding for the structural proteins VP1 and VP2 are typically expressed later in infection, following the accumulation of viral DNA in the nucleus, their expression further supports the permissiveness of a cell to the virus. Consistent with the NS1 mRNA data, quantification of VP1/VP2 mRNA also showed a similar trend in first-trimester CTBs (Fig. 6C). Notably, the substantial increases in NS1 and VP1/VP2 mRNA expression observed in JEG-3 and BeWo cells far exceeded the differences in transfection efficiency, underscoring the higher permissiveness of these CTB-derived cell lines to B19V replication. In contrast, although HTR-8/Svneo cells exhibited the highest transfection efficiency, they did not show a corresponding increase in viral gene expression.

Overall, the findings indicate that the intracellular environment of first-trimester CTBs is significantly more permissive to B19V replication than that of EVTs or CTBs derived from term placentas. As previously suggested, the reduced proliferative capacity of term placental CTBs likely contributes to their lower levels of viral gene expression, highlighting the critical role of cellular proliferation in determining B19V permissiveness. Notably, the first-trimester CTB-derived cell lines, BeWo and JEG-3 exhibited a more permissive intracellular environment than the UT7/Epo cells. However, the absence of globoside expression blocks viral entry, rendering these cells resistant to B19V infection.

### PEI rescues B19V infection in trophoblasts lacking globoside

Globoside plays a pivotal role in the endosomal escape of incoming B19V particles, facilitating their release from endosomes into the cytoplasm, which is a crucial step for viral replication^11^. Several studies have shown that primary trophoblasts, whether isolated from placentas or analyzed in tissue sections, consistently express globoside^18,41,42^. Our findings align with this observation, showing that all trophoblast subtypes tested expressed globoside, with the exception of the two choriocarcinoma-derived BeWo and JEG-3 cell lines, which likely lack it due to tumor-associated changes in glycosphingolipid synthesis^43,44^.

To circumvent the absence of globoside and promote endosomal escape, we employed polyethyleneimine (PEI), a cationic polymer known to enhance endosomal rupture by disrupting endosomal membranes and facilitating the release of internalized particles into the cytoplasm^46^. To evaluate the effect of PEI on endosomal integrity, BeWo and JEG-3 cells were incubated with tetramethylrhodamine-labeled dextrans (3 kDa), which serve as markers of endosomal compartmentalization. In untreated cells with intact endosomes, the rhodamine-labeled dextrans remained confined within discrete vesicles. In contrast, PEI-treated cells (≥1 μM) exhibited diffuse cytoplasmic fluorescence, indicating loss of endosomal integrity (Fig. 7A and B).

**Figure 7.**
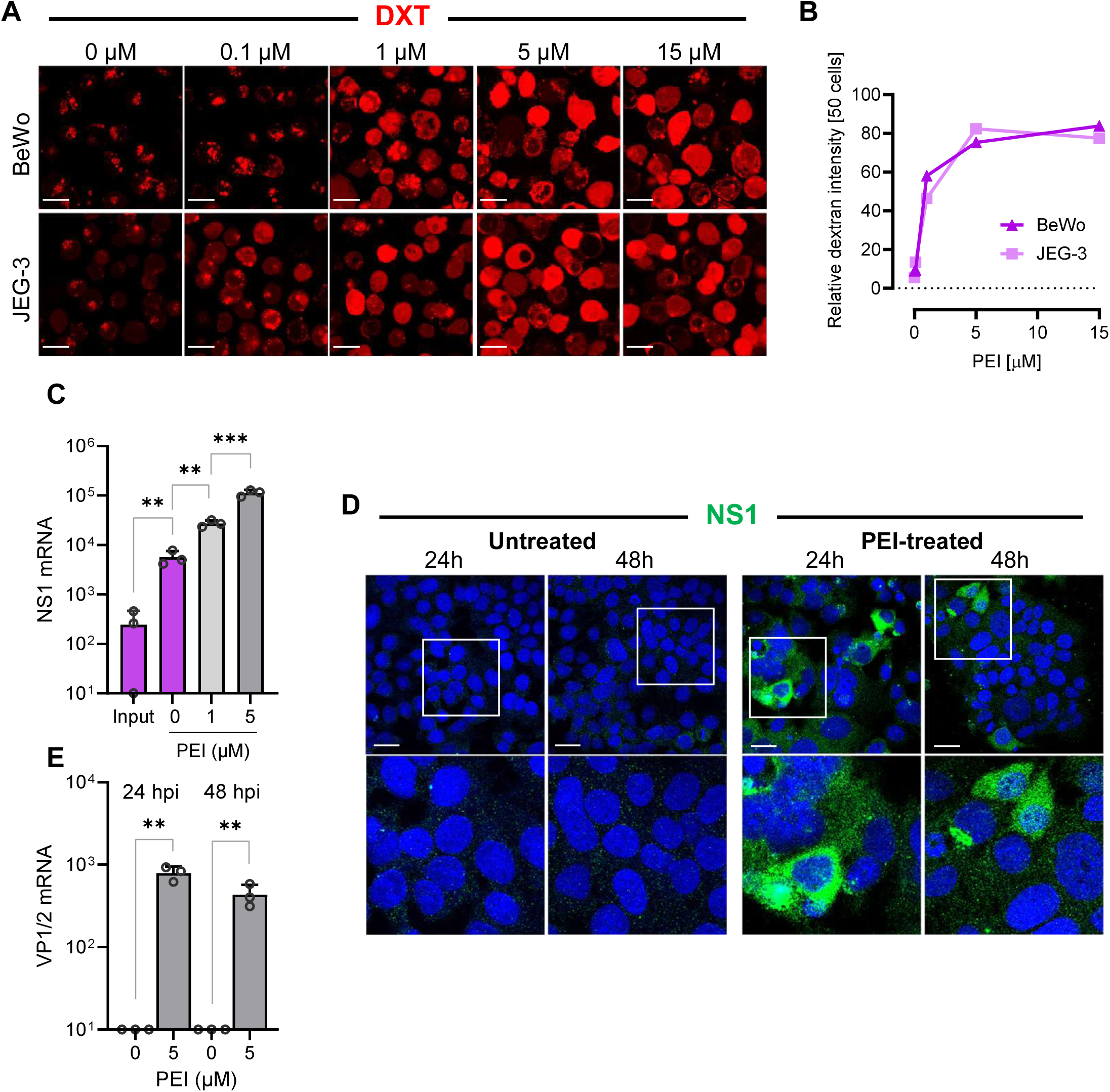
PEI-mediated endosomal escape rescues B19V infection in globoside-deficient trophoblasts. (A) Immunofluorescence images showing intracellular distribution of fluorescent dextrans (DXT: red) in BeWo and JEG-3 cells following treatment with increasing concentrations of PEI. Diffuse cytoplasmic localization of dextrans indicates release from endosomes and disruption of endosomal membranes. Scale bar, 20 μm. (B) Quantification of cells with cytosolic distribution of dextrans in BeWo and JEG-3 cells in response to PEI treatment. Fluorescence intensity was measured in 50 cells per condition using ImageJ. (C) Quantification of NS1 mRNA levels at 24 h post-infection in BeWo cells treated or untreated with PEI. (D) Detection of NS1 protein by immunofluorescence using an NS1-specific antibody (mAb 1424; green) in infected BeWo cells at 24 h and 48 h post-infection, with or without 5 µM PEI treatment. Scale bar, 30 μm. (E) Quantification of structural VP1/VP2 mRNA levels at 24 h and 48 h post-infection in BeWo cells treated or untreated with 5 µM PEI. All results are presented as the mean ± SD of three independent experiments. Statistical significance was calculated using two-sided Student’s t-test. ∗∗*p* < 0.01; ∗∗∗*p* < 0.001.

Cells were infected with B19V in the presence of PEI (1 µM or 5 µM) at 37°C for 30 min. After removing unbound virus and PEI by washing, the infection was allowed to proceed for 24 h, and NS1 mRNA levels were quantified by RT-qPCR. The results revealed a dose-dependent increase in NS1 mRNA levels in PEI-treated cells, indicating enhanced viral replication (Fig. 7C).

To confirm that the transcribed NS1 mRNA was effectively translated into protein, the presence of NS1 protein was examined by immunofluorescence microscopy with an antibody against B19V NS1 (mAb 1424)^47^. NS1 protein accumulation was detected in BeWo cells after 24 h and 48 h post-infection, confirming translation of the NS1 mRNA (Fig. 7D). Given that NS1 is essential for viral DNA replication and the subsequent expression of structural proteins, the expression of VP1/2 mRNA was tested. RT-qPCR confirmed the presence of structural mRNA in PEI-treated BeWo cells, which correlated with the observed NS1 protein expression (Fig. 7E). These results provide compelling evidence that PEI facilitates endosomal escape of incoming viruses, thereby circumventing the lack of globoside in BeWo cells.

### EpoR signaling is not required for B19V infection of trophoblasts

EpoR is a cell surface receptor that binds Epo, a hormone known for its role in erythropoiesis. Although EpoR signaling is not required for virus entry, it has been shown to be essential for B19V replication in EPCs and is thought to be a key factor underlying the pronounced tropism of B19V for human erythroid progenitors^48,49^.

Both Epo and EpoR are expressed by trophoblasts in the human placenta, where EpoR signaling contributes to critical processes such as cell survival, angiogenesis, placental development, and adaptation to hypoxic conditions^38,50^. Although Epo expression has been observed in BeWo cells^51^, quantitative data is not available, and expression in other trophoblast-derived cell lines, such as JEG-3, has not been assessed. To address this gap, we quantified Epo mRNA levels in BeWo and JEG-3 cells under normoxic and hypoxic conditions, using HepG2 and UT7/Epo cells as positive and negative controls, respectively. Epo mRNA was detectable in both trophoblast lines, albeit at substantially lower levels than in HepG2 cells, indicating low, nearly undetectable endogenous Epo expression (Fig. 8A and B). Given the low baseline expression, we examined whether exogenous Epo supplementation could modulate BeWo cell proliferation or survival. However, supplementation with exogenous Epo had no significant impact on BeWo cells, in contrast to UT7/Epo cells, where exogenous Epo is essential for both cell proliferation and viability (Fig. 8C). Transfection of BeWo cells with B19V DNA genomes, either in the presence or absence of exogenous Epo, revealed no substantial differences in NS1 and VP expression, suggesting that EpoR signaling is not essential for B19V infection in trophoblasts (Fig. 8D).

**Figure 8.**
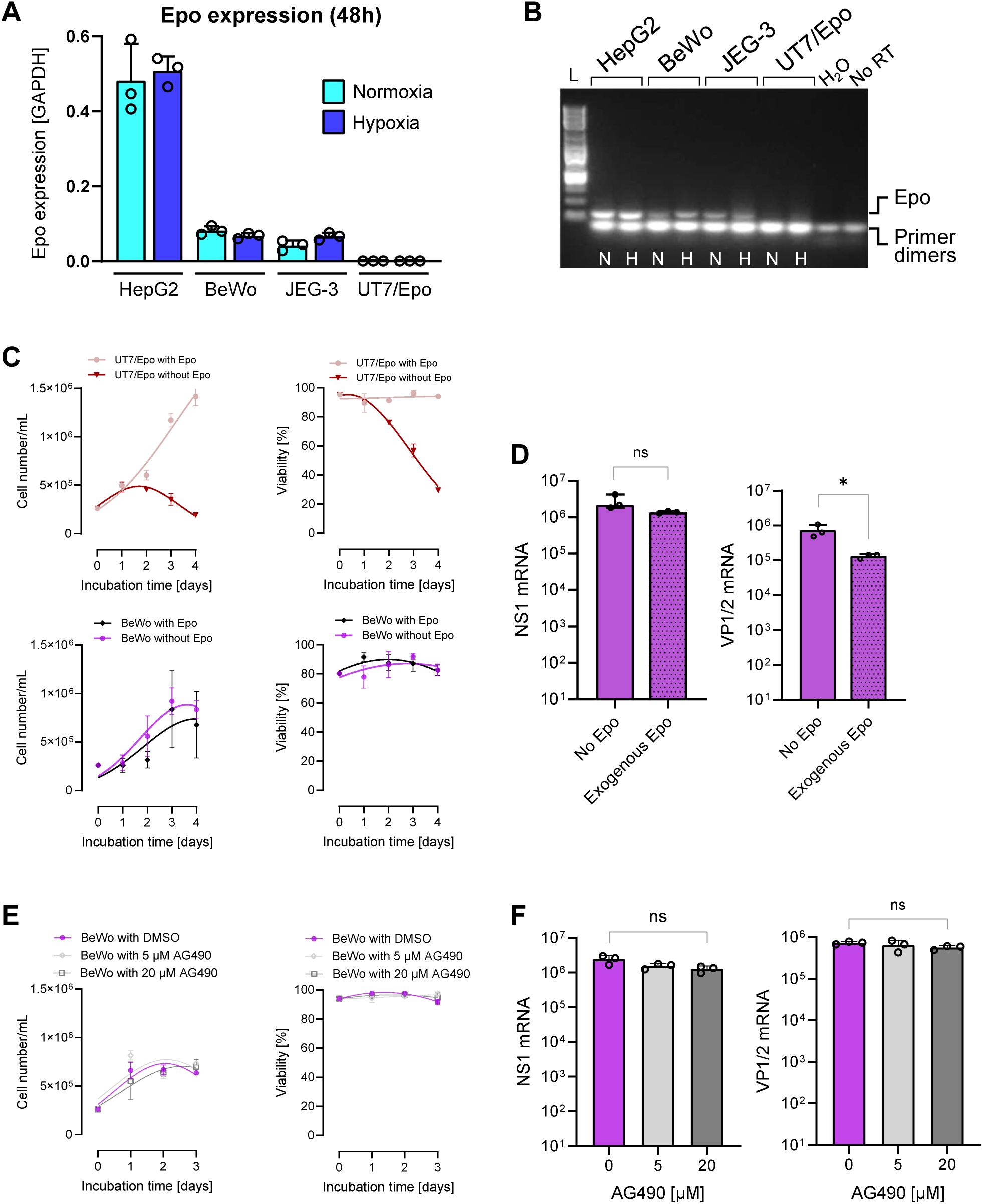
EpoR signaling is dispensable for B19V infection in trophoblasts. (A) Quantification of Epo mRNA levels in BeWo and JEG-3 cells under normoxia (N) and hypoxia (H), with HepG2 and UT7/Epo as positive and negative controls, respectively. (B) Agarose gel electrophoresis of RT-PCR products showing expected amplicon size. L, 1 kb DNA ladder; no RT, no reverse transcriptase. (C) Effect of exogenous Epo supplementation on proliferation and viability of UT7/Epo and BeWo cells. (D) NS1 and VP mRNA levels in BeWo cells transfected with full-length B19V genomes, with or without Epo supplementation. (E) Cell proliferation and viability of BeWo cells treated with 5 or 20 µM AG490. (F) NS1 and VP mRNA expression in BeWo cells treated with AG490 prior to B19V genome transfection.

Both BeWo and JEG-3 cells exhibited low but detectable levels of Epo expression, raising the possibility that even minimal EpoR signaling might be sufficient to support B19V infection. JAK2 is a central mediator of EpoR signaling, activating downstream STAT5 and MEK/ERK pathways, and its inhibition with AG490 has previously been shown to disturb B19V infection in erythroid progenitor cells^48^. AG490 is a tyrphostin compound that blocks JAK2 phosphorylation by competing with ATP^52^. At concentrations below 20 µM, AG490 had no significant effect on the proliferation or viability of BeWo cells (Fig. 8E). To assess the impact of JAK2 inhibition on B19V gene expression, BeWo cells were treated with 5 or 20 µM AG490 prior to transfection with full-length B19V genomes. Subsequent quantification revealed that NS1 and structural VP mRNA levels remained unchanged at both inhibitor concentrations, indicating that JAK2 activity does not influence viral gene expression in these cells (Fig. 8F). Collectively, our findings indicate that B19V infection in trophoblasts does not rely on EpoR signaling, unlike its essential dependence in erythroid progenitor cells.

## Discussion

Human parvovirus B19 primarily infects EPCs in the bone marrow. While well known for causing hematopoietic dysfunction, B19V has gained increasing attention in the context of pregnancy due to its potential to induce severe fetal complications, including fetal anemia, non-immune fetal hydrops, and intrauterine fetal death^3^. These complications are more frequent when infection occurs during the first trimester, coinciding with a developmental window characterized by heightened placental plasticity and rapid trophoblast proliferation^15–17^. Despite the recognition of these risks, knowledge of the mechanism of B19V transmission to the fetus remains poorly understood. As no targeted antiviral therapies or vaccines are currently available, the management of fetal B19V infection relies primarily on monitoring for signs of fetal anemia and, when necessary, performing intrauterine blood transfusions^53^.

B19V infection is primarily mediated by VP1uR, which has a highly restricted expression pattern limited to EPCs in the bone marrow^6–8^. This restricted expression profile underlies the pronounced erythroid tropism characteristic of B19V. In this study, we show that B19V also infects placental trophoblasts via the same receptor-mediated entry pathway, thereby expanding the known cellular tropism of the virus beyond the erythroid lineage. In contrast to CTBs and STBs, EVTs, which represent a terminally differentiated lineage derived from CTBs, do not express VP1uR and, correspondingly, do not support B19V uptake and infection. This mirrors our observations during erythropoiesis, where VP1uR is present in early erythroblasts but downregulated in terminally differentiated erythroid cells^8^. These observations suggest that VP1uR may be a differentiation-associated receptor, expressed at intermediate stages of cellular maturation. Of note, the lack of detectable VP1uR in the EVT model HTR/SVneo does not necessarily imply that placental EVTs lack VP1uR expression. This finding requires validation using additional EVT cell models and, ideally, confirmation in placental tissue.

Although the tyrosine kinase AXL has recently been proposed as a receptor for VP1u in erythroid cells^34^, its broad expression across various human tissues^54^ does not align with the strict erythroid tropism of B19V. Supporting this, our data show that EVTs, which express AXL but lack VP1uR, are not susceptible to B19V infection. In contrast, BeWo and JEG-3, which express VP1uR but not AXL, do support viral uptake. Together, these findings indicate that AXL is not required for B19V entry into placental trophoblasts.

Previous work demonstrated that the virus binds globoside only under acidic conditions^9^, preventing unwanted interactions in non-permissive cells where globoside is widely expressed. This pH-dependent binding occurs in the acidic environment of the nasal mucosa, allowing the virus to attach to epithelial surfaces and undergo transcytosis^10^. Beyond the nasal mucosa, globoside is essential for endosomal escape of B19V following VP1uR-mediated endocytosis in EPCs. In globoside knockout UT7/Epo cells, incoming viruses remain trapped in endosomes, preventing infection. In these cells, infectivity is restored by PEI-induced endosomal rupture^11^. Numerous studies have shown that globoside is expressed in the human placenta^18,41,42,55^, which we confirmed both in trophoblast cell lines and in cryosections from term placenta. Although first-trimester placenta sections were not available, these results strongly suggest that globoside is widely expressed in the human placenta. The only exception in our analyses were the choriocarcinoma-derived cell lines BeWo and JEG-3. The lack of globoside expression is likely due to the disrupted glycosylation pathways in these cancer-derived cell lines, which are known to exhibit altered glycosyltransferase expression compared to the normal cells from which they derive^43,44^. Although BeWo and JEG-3 cells express VP1uR and support viral internalization, productive replication is blocked, indicating that globoside is essential for B19V infection in trophoblasts. As in globoside KO UT7/Epo cells^11^, PEI-mediated endosomal rupture restored viral replication, confirming that globoside is required for endosomal escape of B19V in trophoblasts, underscoring the virus-globoside interaction as a potential antiviral target. These results also emphasize that, although BeWo and JEG-3 cells are widely used to study trophoblast biology, their lack of globoside expression renders them unsuitable for modeling B19V infection.

Our findings indicate that both CTB and STB populations are susceptible to B19V. However, the markedly higher permissiveness of first-trimester CTBs, which are highly proliferative compared to the low-proliferating CTBs from term placentas suggests that cellular proliferation is a key determinant of productive infection. As trophoblasts transition toward a more differentiated, functionally specialized state with reduced proliferative capacity in later gestation, their ability to support B19V replication appears to diminish. This differential permissiveness supports a model in which the developmental stage of placental trophoblasts shapes their permissiveness to the infection and may explain the higher frequency of fetal infection during early pregnancy. Supporting this, BeWo and JEG-3 cells, both derived from first-trimester CTBs, produce significantly higher levels of NS1 mRNA upon transfection compared to term placenta-derived hPTC^CTB^ cells. However, HTR-8/SVneo cells, which are EVTs from early placenta and exhibit similar proliferation rates to BeWo and JEG-3, generate lower NS1 mRNA levels. This suggests that, beyond proliferation, additional intrinsic intracellular factors contribute to the elevated permissiveness of early CTBs compared to EVTs.

Together, these observations suggest that B19V permissiveness in trophoblasts is governed by a combination of proliferative capacity and cell-intrinsic factors. Building on this, we explored whether signaling pathways known to be essential for the infection in erythroid progenitors also contribute to infection in placental cells. Interestingly, the fact that EpoR signaling is not required for B19V infection in trophoblasts stands in contrast to findings in erythroid cells, where Epo-induced JAK2 activation is critical for productive infection, a dependence that contributes to the strict erythroid tropism of the virus^48^. Although the underlying mechanisms remain to be elucidated, this points to the existence of alternative, Epo-independent pathways that enable virus replication in non-erythroid cells, thereby broadening our understanding of the cellular tropism and adaptability of B19V.

In summary, this study demonstrates that B19V can enter and infect placental trophoblasts, expanding its recognized strict tropism beyond the erythroid lineage to include early villous trophoblasts. The combined presence of VP1uR, globoside, and high proliferative capacity underlies the heightened susceptibility of the early placenta and positions first-trimester villous trophoblasts as the most relevant models for elucidating infection mechanisms and guiding the development of antiviral strategies to prevent vertical transmission.

### Limitations of the study

This study used trophoblast models, which, while useful for identifying key factors mediating B19V entry and permissiveness, do not fully capture the complexity of the placental environment. Immune and endothelial cells, as well as broader tissue interactions, were not represented. The work focused on entry and replication, without assessing downstream effects such as immune responses or barrier disruption. Additional trophoblast models should be examined to more accurately delineate B19V tropism within the placenta. Nonetheless, by defining the cellular determinants of B19V infection, this study provides a mechanistic basis for future investigations using first-trimester villous trophoblasts and more physiologically relevant models, such as trophoblasts organoids or multi-cell-type 3D cultures.

## Resource availability

### Lead contact

Further information and requests for resources and reagents should be directed to the lead contact, Carlos Ros.

### Materials availability

All unique reagents generated in this study are available from the lead contact upon request, subject to a materials transfer agreement.

### Data and code availability

All study data are included in the main text or the supporting information.

## Acknowledgments

We thank Tina Bürki (Swiss Federal Laboratories for Material Science and Technology, EMPA) for kindly providing BeWo b30 Aberdeen choriocarcinoma cells. We are grateful to Christiane Albrecht and Stefan Rudloff (University of Bern) for providing JEG-3 and HTR-8/SVneo cells, respectively, and Ramkumar Menon (University of Texas) for the hPTC^CTB^ cells. This study was supported by a grant from the Swiss National Science Foundation (grant 320030_207850 to C.S.).

## Author contributions

Conceptualization, C.S., and C.R.; methodology, C.S., M.K., J.B., and A.F.; data curation and formal analysis, C.S., M.K., J.B., and C.R.; resources, C.S., D.B., M.A., and C.R.; writing - review & editing, C.S., and C.R.; manuscript revision, all authors; funding acquisition, project administration, and supervision, C.R.

## STAR★Methods

### Key resources table

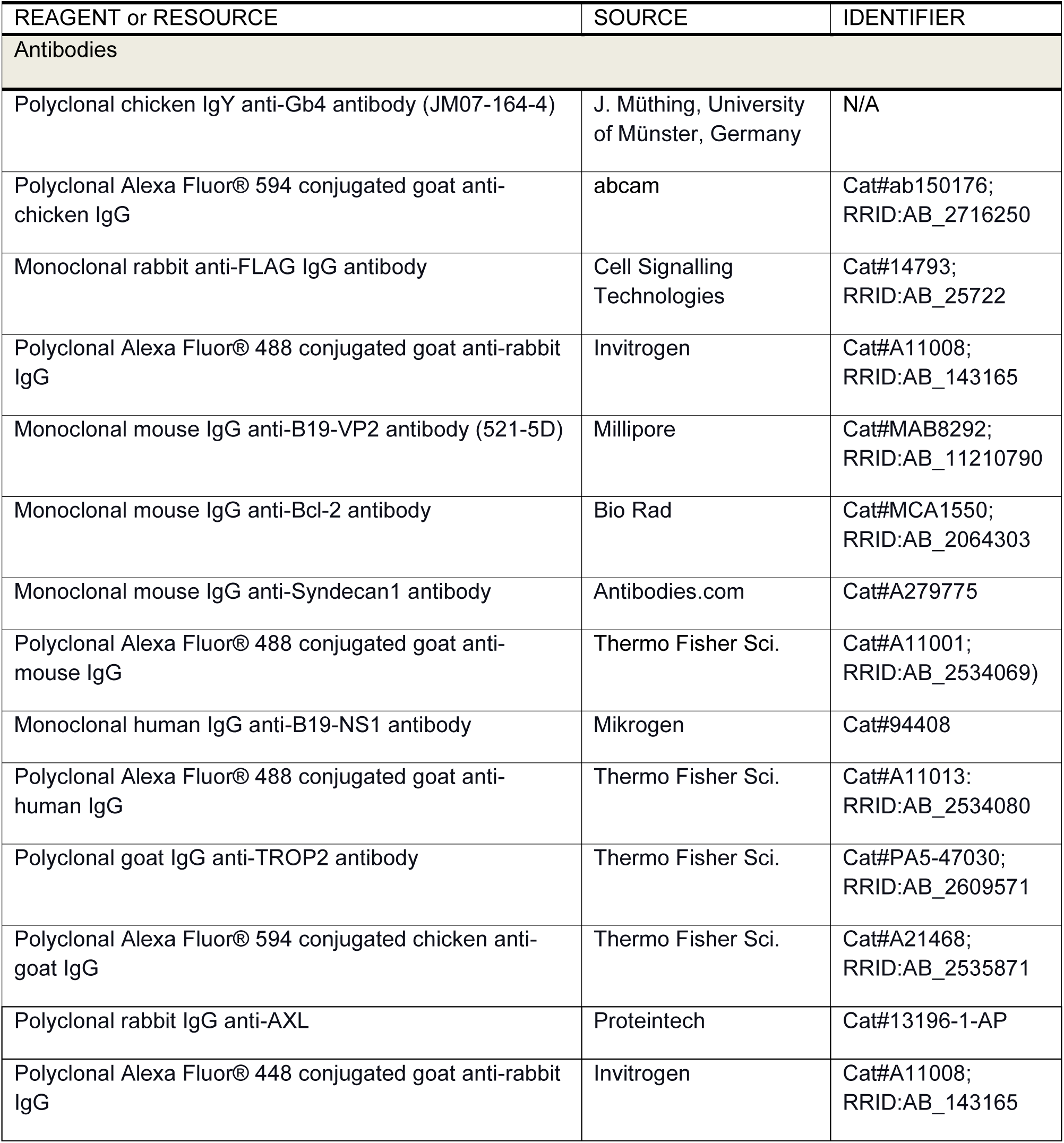
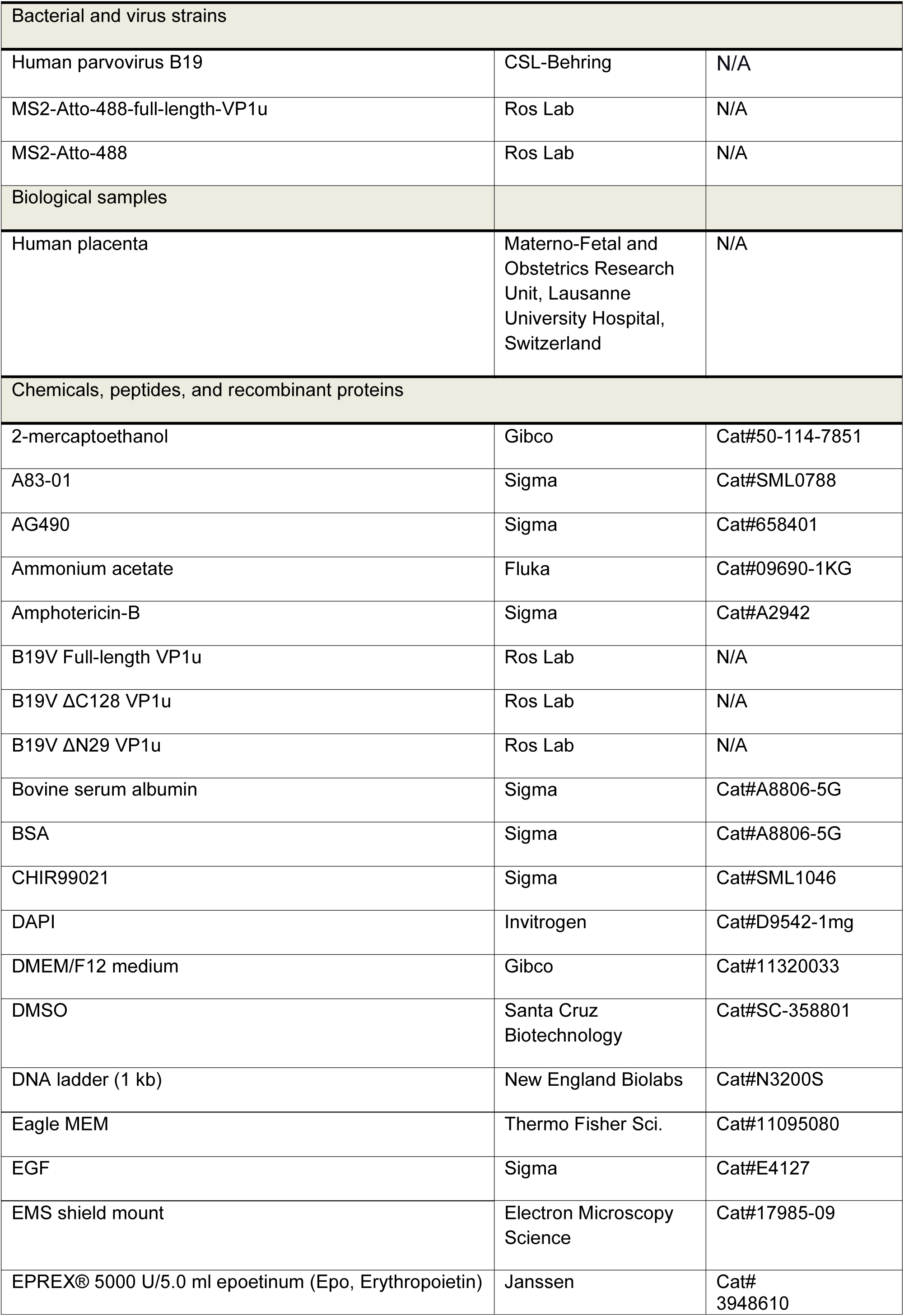
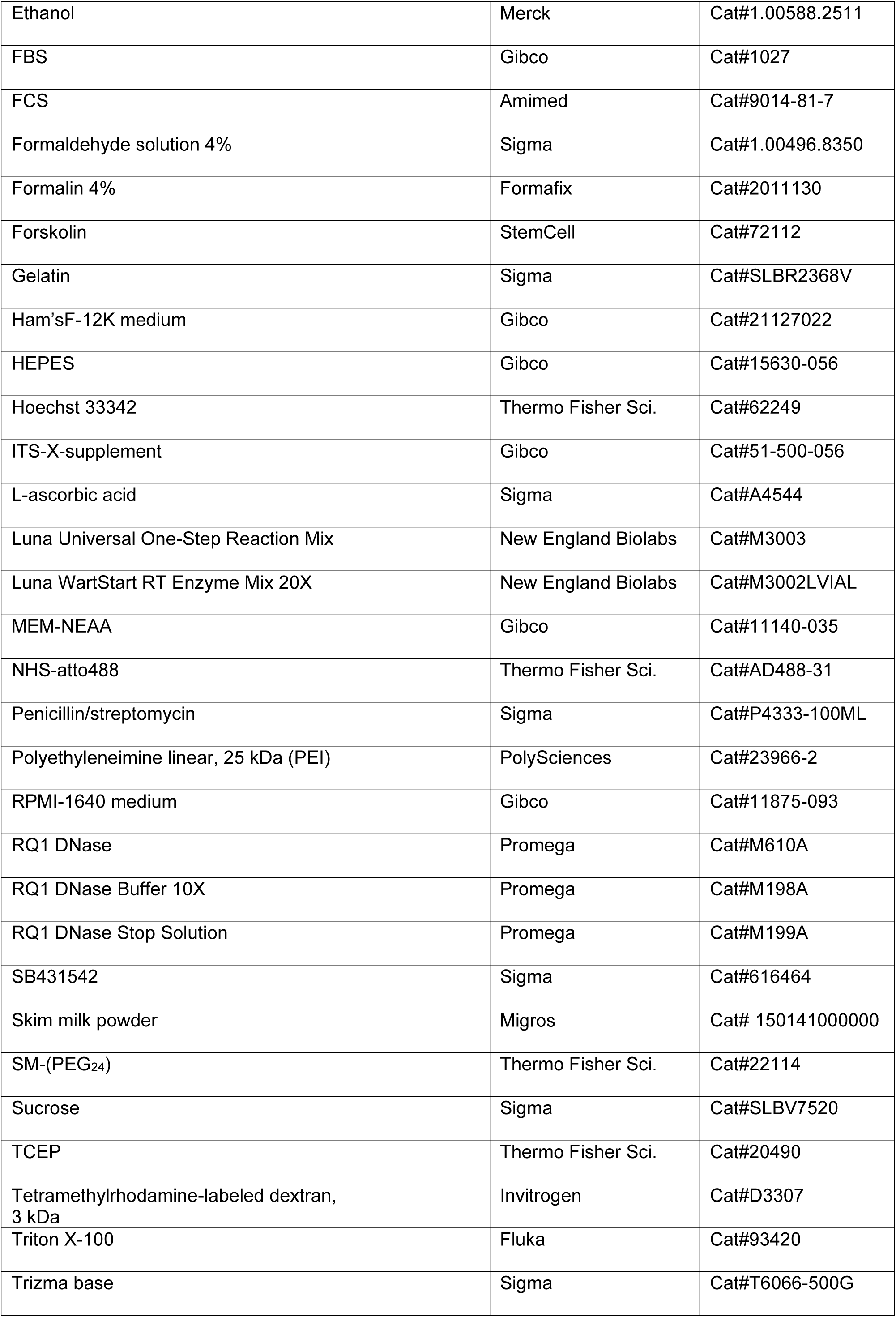
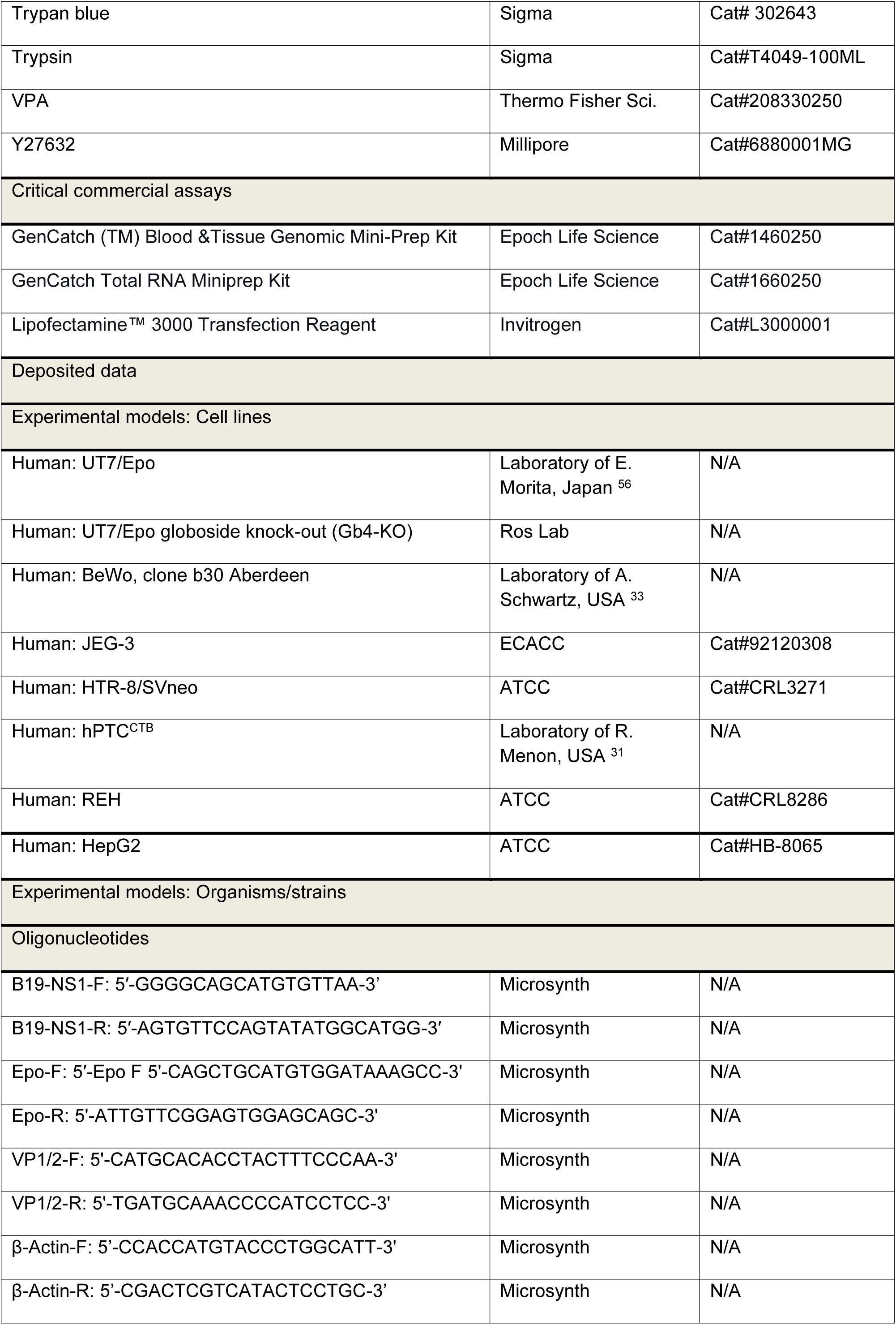
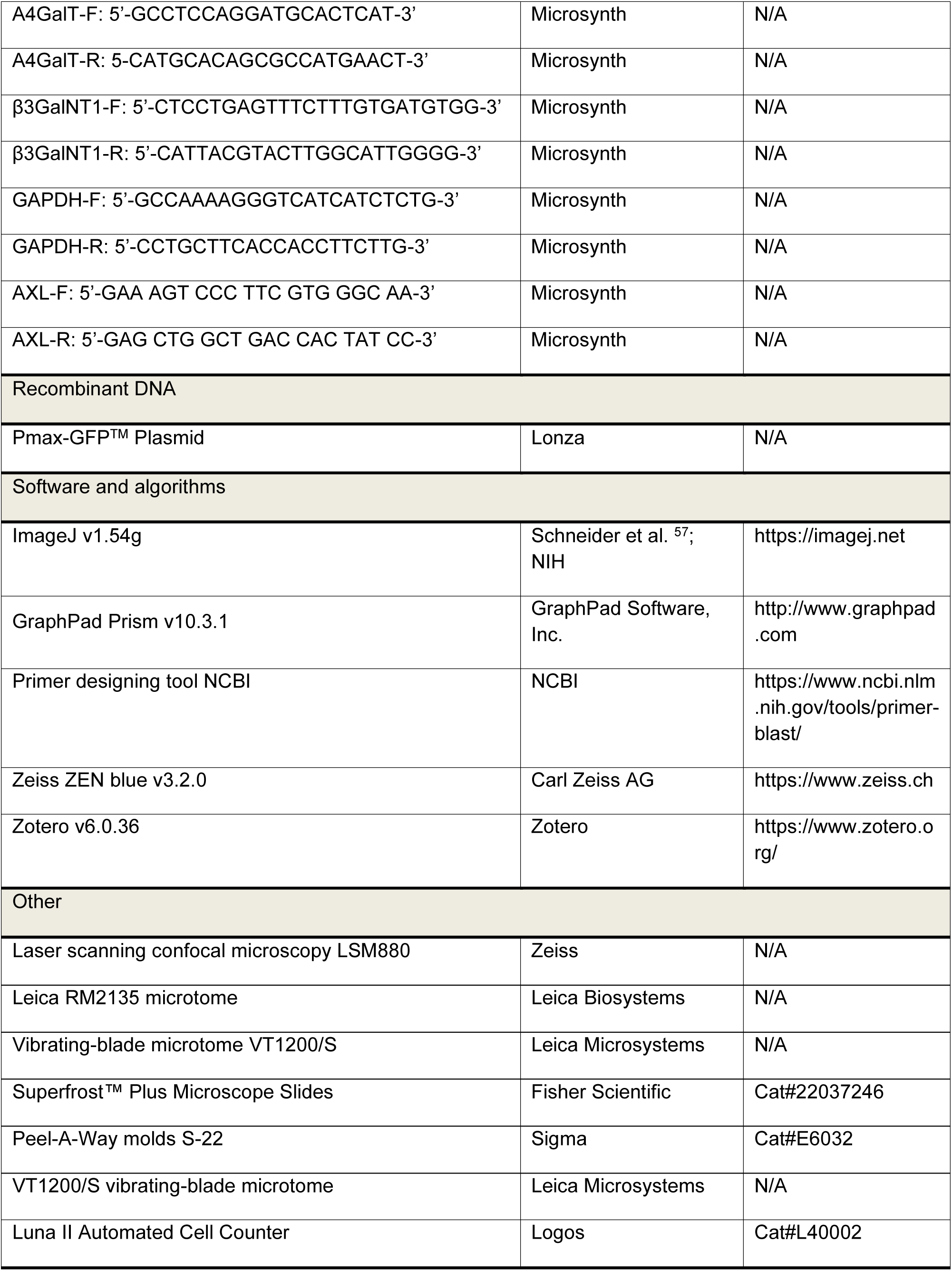

## Method details

### Cells

UT7/Epo cells, provided by E. Morita (Tohoku University School of Medicine, Japan) were cultured in Eagle’s Minimum Essential Medium (MEM, Thermo Fisher Scientific, Waltham, MA, USA) with 5% FCS and 2 U/mL Epo. UT7/Epo Gb4-KO cells were generated by co-transfecting a CRISPR/Cas9 GFP plasmid targeting β-1,3-Gal-T3 with an HDR RFP plasmid, followed by bulk and single-cell FACS sorting and confirmed by RT-qPCR as previously described^35^. The UT7/Epo globoside-KO cells are cultured in the same manner as UT7/Epo cells. The BeWo b30 Aberdeen choriocarcinoma cells were obtained from T. Bürki (Swiss Federal Laboratories for Material Science and Technology; EMPA, St. Gallen, Switzerland) and cultured in Ham’s F-12K medium (Gibco, Waltham, MA, USA) containing 10% FCS and 50 U/mL of penicillin/streptomycin. BeWo b30 Aberdeen cells were differentiated into syncytiotrophoblasts with 10 μM of forskolin (72112, STEMCELL Technologies Switzerland GmbH, Basel, Switzerland) at 37°C for 72 h. The choriocarcinoma JEG-3 cells were obtained from C. Albrecht (Institute of Biochemistry and Molecular Medicine, University of Bern, Switzerland). The hepatocarcinoma cell line HepG2 was purchased form ATCC. JEG-3 and HepG2 cells were cultured in Eagle’s MEM with 5% FCS and 50 U/mL penicillin/streptomycin. The extravillous trophoblast cell line HTR-8/SVneo was obtained from S. Rudloff (Department of Nephrology and Hypertension, University of Bern, Switzerland). The cells were grown in RPMI1640 medium (Gibco) containing 5% FCS and 50 U/mL penicillin/streptomycin. The REH cells, a B-cell precursor leukemia cell line, were cultured in RPMI 1640 medium containing 10% FCS and 50 U/mL penicillin/streptomycin. The aforementioned cells were routinely maintained under standard conditions at 37°C and 5% CO_2_. The hPTC^CTB^ cells, an immortalized term placenta-derived trophoblast cell line, were a gift from R. Menon (Department of Obstetrics and Gynecology, Maternal-Fetal Medicine, Perinatal Research Division, University of Texas). The cells were grown in DMEM:F12 medium supplemented with 0.2% FCS, 0.3% BSA, 1% amphotericin-B (AmpB), 1% ITS-X-supplement, 1% penicillin/streptomycin, 50 ng/mL EGF, 0.5 μM A83-01, 1 μM SB431542, 1.5 μg/mL L-ascorbic acid, 2 μM CHIR99021, 5 μM Y27632, 0.8 mM VPA, and 0.1 mM 2-mercaptoethanol and cultured at 37°C, 5% CO_2_, and 5% O_2_. The medium was replenished every second day. All cells were used at low passage numbers (P5–25) to minimize genetic drift and phenotypic changes.

### Cryosections of human placenta

Placental tissue was obtained directly after birth from term pregnancies (37-42 weeks) through caesarean section in patients aged 25-35 years by cutting full-thickness biopsies (∼5 cm). Written informed consent was obtained from all the patients, and the study protocol agrees with the local Ethics Committee of the Canton of Vaud, Switzerland.

Residual blood was removed by washing the biopsies with ice-cold PBS (Gibco) supplemented with 100 U/mL penicillin and 100 µg/mL streptomycin. The tissue was trimmed to approximately 1 cm³, embedded in 1% of low melting point agarose (Promega, Madison, WI, USA) in Peel-A-Way molds S-22 (Sigma, St. Louis, MO, USA) and kept on ice. Using a VT1200/S vibrating-blade microtome (Leica Microsystems), sections of 700 μm thickness were obtained with the following settings: speed 0.12-0.26 mm/s, amplitude 3 mm, angle 18°. The human placenta slices were transferred to 6 well plates (TPP) containing 3 mL DMEM GlutaMax (32430027, Gibco), 10% fetal bovine serum (10270, Gibco), 10 mM HEPES (1530056, Gibco), 1% Glutamine, 1% MEM-NEAA, 100 units/ml of penicillin and 100 μg/ml streptomycin. The cultures were maintained for 2 days at 37°C, 5% CO_2_ prior infection and medium was changed every 24 h.

The specimens were fixed in 4% formalin (Formafix) over night at 4°C. For cryopreservation, the tissue was first incubated in 30% sucrose/PBS overnight, embedded in 7.5% gelatine/10% sucrose at 37°C for 45 min, and polymerized at room temperature. Samples were frozen in a dry ice/100% ethanol bath and stored at -80°C. Cryosections of 10 μm were cut using a Leica cryostat and mounted on Superfrost Plus glass slides (Thermo Fisher Scientific) as previously described^58^.

### Viruses

Native B19V was obtained from de-identified plasma samples of infected, seronegative individuals, confirmed by virus-specific serology (CSL Behring, Bern, Switzerland). Infected plasma was thawed and clarified by centrifugation at 4,000 rpm for 10 minutes at 4°C. Viral genomes were quantified by quantitative PCR (qPCR) using Luna Universal One-Step Reaction Mix (M3003, New England Biolabs, Ipswich, MA, USA) with primers specific for the NS1-coding region as mentioned below. Plasmids containing the complete B19V genome were used as external standards in 10-fold serial dilutions.

### Expression and purification of recombinant VP1u and MS2 VLPs

*Escherichia coli* (*E. coli*) BL21(DE3) containing pT7-FLAG-MAT-Tag-2 vectors encoding for full-length or truncated (ΔN29, ΔC128) VP1u variants of B19V were grown in LB broth medium and induced with ITPG for protein expression. Proteins were harvested and purified by Ni-NTA affinity chromatography as described previously^6^. For click-chemistry, recombinant VP1u was reduced with 5 mM TCEP.

MS2-bacteriophage virus-like particles were expressed using *E. coli* BL21(DE3). MS2-VLPs were purified by ultracentrifugation through a 20% sucrose cushion as described elsewhere^8^.

### Conjugation of VP1u to fluorescently labeled MS2 VLPs by click chemistry

MS2 VLPs were purified and labelled with NHS-Atto 488 (Atto-Tec, Siegen, Germany) using a 40-fold molar excess of dye. The reaction was quenched, and labelled VLPs were recovered by centrifugation through a 20% sucrose cushion to remove unreacted dye. Subsequently, MS2 VLPs were incubated with 500-fold molar excess of maleimide-PEG24-NHS (22114, Thermo Fisher Scientific) for 1 h. Excess crosslinker was eliminated using a 40 kDa MWCO desalting column. The activated VLPs were then conjugated to reduced recombinant VP1u protein and further purified by a second 20% sucrose cushion^8^.

### Drugs and reagents

Forskolin (STEMCELL Technologies Switzerland GmbH, Basel, Switzerland) was dissolved in DMSO at 10 mM. Linear 25 kDa polyethyleneimine (PEI; PolySciences, Warrington, PA, USA) was dissolved in water at 1 mM. A 3 kDa tetramethylrhodamine-labeled dextran (Invitrogen, Carlsbad, CA, USA) was dissolved in water at 20 mg/mL. CHIR99021 (SML1046), A83-01 (SML0788), SB431542 (616464), L-ascorbic acid (A4544), EGF (E4127), Penicillin/Streptomycin (P4333-100ML), and Amphotericin-B (A2942) were obtained from Sigma. CHIR99021 and A83-01 were dissolved in DMSO; L-ascorbic acid was dissolved in water; and EGF was dissolved in 0.85% NaCl. The reagent Y27632 was obtained from Millipore (6880001MG, Darmstadt, Germany) and dissolved in water. The ITS-X-supplement (51-500-056) and 2-mercaptoethanol (50-114-7851) were obtained from Gibco. BSA and the VPA were obtained from Thermo Fisher Scientific. AG490 (658401) was purchased from Sigma and dissolved in DMSO.

### Binding and internalization assays

Binding of recombinant VP1u (100 ng) and B19V (10^5^ geq/cell) was performed at 4°C for 1 h in UT7/Epo, BeWo, JEG-3, hPTC^CTB^, and REH cells resuspended in PBS (pH 7.2). Cells were washed with ice-cold PBS and then prepared for internalization, DNA extraction, or fixed for immunofluorescence. DNA was isolated using the GenCatch Plus Genomic DNA Miniprep Kit (1660250, Epoch Life Science, Missouri City, TX, USA). Internalization of recombinant VP1u (100 ng), B19V (10^5^ geq/cell), and MS2-VP1u (10^6^ geq/cell) was conducted in UT7/Epo, BeWo, JEG-3, hPTC^CTB^, and REH cells at 37°C for 30 min. Cells were then washed and either fixed for immunofluorescence or processed for DNA extraction, as described above. Viral DNA was analysed by qPCR with the following primers: B19-NS1-F: 5′-GGGGCAGCATGTGTTAA-3′, B19-NS1-R: 5′-AGTGTTCCAGTATATGGCATGG-3′.

### Immunofluorescence

For surface detection of globoside, cells of interest were blocked with 1% bovine-serum albumin prior to incubation with a polyclonal chicken IgY anti-Gb4 antibody (JM07-164-4, J. Müthing, University of Münster, Germany) at 4°C for 30 minutes. Cells were washed multiple times with ice-cold PBS before fixation with 4% formaldehyde for 10 min, washed with PBS, quenched with 1 M Tris-HCl (pH 8.5) as previously described^10^. The antibody targeting globoside was detected with a polyclonal goat anti-chicken IgG antibody conjugated to Alexa Fluor® 594 (ab150176, abcam, Cambridge, UK). Detection of VP1uR was assessed in cells of interest by incubating recombinant B19 VP1u at 37°C for 30 min in the presence of a monoclonal rabbit anti-FLAG IgG antibody (14793S, Cell Signalling Technologies, Danvers, MA, USA). The cells were washed several times with ice-cold PBS before fixation in a 1:1 mixture of methanol: acetone at -20°C for 4 min. The FLAG antibody was detected with a polyclonal Alexa Fluor® 488 conjugated goat anti-rabbit IgG (A11008, Invitrogen). B19V was detected using a monoclonal mouse IgG anti-B19-VP2 antibody (521-5D; MAB8292, Millipore, Burlington, MA, USA) and a secondary polyclonal goat anti-mouse IgG conjugated to Alexa Fluor® 488 (A11001, Invitrogen). The nonstructural protein NS1 was detected using a monoclonal human IgG anti-B19-NS1 antibody (1424; 94408, Mikrogen, Neuried, Germany) and a polyclonal goat anti-human IgG antibody conjugated to Alexa Fluor® 488 (A11013, Invitrogen). AXL was detected using a polyclonal rabbit IgG anti-AXL antibody (13196-1-AP, Proteintech, Manchester, UK) and a polyclonal goat anti-rabbit IgG antibody conjugated to Alexa Fluor® 488 (A11008; Invitrogen). The samples were mounted with EMS shield mount (17985-09, Electron Microscopy Science, Hatfield, PA, USA) containing 0.2 ng/mL DAPI. The samples were visualized with a 63× oil immersion objective by laser scanning confocal microscopy (LSM880, Zeiss, Oberkochen, Germany).

For immunofluorescence analysis of FFPE slices, gelatine was removed, formalin was quenched with 1 M Tris-HCl, pH 8, tissue was permeabilized with 0.2% Triton-X-100 and blocked with 5% milk/PBS. Cytotrophoblasts were labelled using a polyclonal goat IgG anti-TROP-2 antibody (1:100, PA5-47030, Thermo Fisher Scientific) followed by a polyclonal chicken anti-goat IgG antibody conjugated to Alexa Fluor® 594 (A21468, Thermo Fisher Scientific). Syncytiotrophoblasts were detected using a monoclonal mouse IgG anti-Bcl-2 antibody (1:50, MCA1550, Bio-Rad, Hercules, CA, USA), and the apical surface of syncytiotrophoblasts was detected using a monoclonal mouse IgG anti-Syndecan-1 antibody (1:100, A279775, Antibodies.com, Cambridge, UK) and a polyclonal goat anti-mouse IgG antibody conjugated to Alexa Fluor® 488 (A11001, Thermo Fisher Scientific). All antibodies were diluted in 2% milk/PBS and carried out at 4°C, after which specimens were treated with 2 mM CuSO_4_/50 mM NH_4_. Specimen was mounted with EMS shield mount as described above and visualized using a 63× oil immersion objective by LSM880.

### Infectivity assays

Cells were infected with B19V (10^5^ geq/cell) at 37°C for 30 min before cells were washed with PBS and then either incubated further at 37°C or processed for RNA extraction using the GenCatch Total RNA Miniprep Kit (1660250, Epoch Life Science). Extracted RNA was DNase I-treated at 37°C for 1 h before NS1 mRNA was quantified by RT-qPCR using the primers as previously specified. When PEI was used for infection, cells were inoculated with PEI (15 μM) simultaneously and washed with PBS at specific times post-infection. RNA was extracted and analysed by RT-qPCR. Following primers were used: NS1-F: 5′-GGGGCAGCATGTGTTAA-3′, NS1-R: 5′-AGTGTTCCAGTATATGGCATGG-3′, VP1/2-F: 5’-CATGCACACCTACTTTCCCAA-3’, and VP1/2-R: 5’-TGATGCAAACCCCATCCTCC-3’. Transfection of B19V DNA was carried out with Lipofectamine™ 3000 Transfection Reagent (L3000001, Invitrogen) following the manufacturer’s protocol closely.

### Endosomal integrity

To assess the effect of PEI on endosomal integrity, cells were incubated with increasing concentrations of PEI and low-molecular weight tetramethylrhodamine-labeled dextran (3 kDa) at 37°C for 30 min and washed several times with PBS. Live-cell images were taken using the LSM880 microscope. Fluorescence intensity was measured in 50 cells per condition using ImageJ (National Institutes of Health, Bethesda, MD, USA).

### Transfection

For transfection, cells between passages 5–25 were seeded in 24-well plates using standard growth medium and transfected the following day with 100 ng of purified B19V DNA. Transfection was carried out with Lipofectamine™ 3000 Transfection Reagent (L3000001, Invitrogen) according to the manufacturer’s protocol. Two days post-transfection, cells were washed several times with PBS and DNA was extracted with Epoch Kit as described above.

To analyse the transfection efficiency, cells were transfected with 100 ng GPF-expressing plasmid DNA (Lonza, Basel, Switzerland) using Lipofectamine™ 3000. The nuclei were stained with Hoechst (1 µg/mL) at 37°C for 10 min for live-cell imaging. Images were captured by LSM880 confocal microscopy. The proportion of transfected cells was quantified using ImageJ by counting GFP-positive cells relative to the total number of Hoechst-stained nuclei. Therefore, channels corresponding to DAPI and GFP were separated, and the intensity thresholds were adjusted to a range of 100-255. Cell segmentation was performed by converting images to binary format and touching objects were separated. Particle analysis was conducted with a size range of 0.0005 cm² to infinity and circularity set from 0.00 to 1.00. The number of GFP-positive (transfected) and DAPI-positive (total) cells was quantified. Transfection efficiency was calculated as the percentage of GFP-positive cells relative to the total number of cells.

### Cell proliferation assays

Cell proliferation was monitored over four days. Cells were seeded in standard growth medium in 24-well plates and incubated for zero to four days. At each timepoint, cells were washed with PBS, lysed in the well and genomic DNA was extracted using the Epoch DNA Extraction Kit as mentioned above. The β-actin gene was quantified by qPCR using the following primers: β-Actin-F: 5’-CCACCATGTACCCTGGCATT-3’, β-Actin-R: 5’-CGACTCGTCATACTCCTGC-3’ and displayed as the percentual increase over day 0.

Cell number and viability were monitored over 3-4 days. Cells were either left untreated or treated with 2 U/mL Epo or 20 μM AG490. At each time point, adherent cells were detached using Accutase, resuspended in standard growth medium to the desired final volume, and mixed 1:1 (v/v) with 0.2% trypan blue. Total cell counts and viability were then measured using a LUNA™ automated cell counter (Logos Biosystems, Anyang, South Korea).

### Detection of glycosyltransferase mRNA expression

The expression of glycosyltransferase mRNAs involved in the synthesis of Gb3 and Gb4 was assessed by extracting total RNA from the cells of interest followed by DNase-I treatment. The mRNA of the 4-α-galactosyltransferase (A4GalT) was quantified by RT-qPCR using the primers A4GalT-F: 5’-GCCTCCAGGATGCACTCAT-3’ and A4GalT-R: 5-CATGCACAGCGCCATGAACT-3’. This enzyme adds a galactose to lactosylceramide, creating trihexosylceramide (Gb3). The mRNA of the 3-β-N-acetylgalactosaminyltransferase (β3GalNT1) was quantified using the primers β3GalNT1-F: 5’-CTCCTGAGTTTCTTTGTGAT GTGG-3’ and β3GalNT1-R: 5’-CATTACGTACTTGGCATTGGGG-3’. This enzyme initiates the final enzymatic step that adds the N-acetylgalactosamine to Gb3, producing Gb4. GAPDH quantification was used for data normalization in qPCR experiments using the following primers: GAPDH-F: 5’-GCCAAAAGGGTCATCATCTCTG-3’ and GAPDH-R: 5’-CCTGCTTCACCACCTTCTTG-3’.

### Quantification and statistical analysis

Statistical analyses were carried out using GraphPad Prism version 10.3.1 (GraphPad Software, Boston, Massachusetts USA). Details about the statistical method used are mentioned in the legend section of the respective figures. Error bars indicate SD as mentioned in the respective figure legends. In all cases, a p value <0.05 is considered significant.

