## supplementary figures for "Cellular determinants of parvovirus B19 infection in the human placenta"

### Supplemental information

**Figure S1. Expression of AXL in trophoblasts does not correlate with VP1uR expression and B19V uptake.**

(A) Relative expression of AXL mRNA in UT7/Epo and in different trophoblast subtypes. mRNA levels were measured by RT-qPCR and normalized to GAPDH expression.

(B) RT-PCR amplicons from BeWo and UT7/Epo cells were visualized by agarose gel electrophoresis. GAPDH, loading control; No RT, no reverse transcriptase; L, 1 kb DNA ladder.

(C) Immunostaining of UT7/Epo and trophoblasts with an anti-AXL antibody (green). DAPI (Blue). Scale bar, 20 µm.

All results are presented as the mean ± SD of three independent experiments.

**Figure S2. Lack of globoside synthase (Gb4) expression in BeWo cells under various conditions.**

(A) Relative mRNA expression of Gb3 and Gb4 in BeWo cells treated or untreated with forskolin (FSK), normalized to GAPDH, to assess the effect of syncytiotrophoblast differentiation on Gb3/Gb4 expression.

(B) Relative mRNA expression of Gb3 and Gb4 in BeWo cells exposed to normoxia, normoxia plus Epo, hypoxia, or hypoxia plus Epo, normalized to GAPDH, to assess the effect of oxygen concentration and Epo on Gb3/Gb4 expression.

(C) Agarose gel electrophoresis of RT-PCR products corresponding to Gb3, Gb4, and GAPDH in BeWo cells under the conditions described in (B), and in UT7/Epo cells incubated with or without Epo under normoxia. Each panel displays the PCR products for Gb3, Gb4, or GAPDH as indicated. No RT, no reverse transcriptase; 1 kb DNA ladder.

All results are presented as the mean ± SD of three independent experiments. Statistical significance was calculated using two-sided Student’s t-test. ∗∗$p < 0.05$; ns, non-significant.

**Figure S3. Expression of globoside in term placenta.**

(A) Trophoblast markers BCL-2, Trop-2 (both in green), and CD138 (red), detected in term placenta cryosections. DAPI (Blue). Scale bar, 40 µm.

(B) Globoside (red), detected in term placenta cryosections. DAPI (Blue). Scale bar, 40 µm.

**Figure S4. Time-dependent increase in β-actin mRNA levels as a proxy for cell growth in trophoblast subtypes.**

Relative β-actin DNA levels were quantified by RT-qPCR in different trophoblast subtypes, UT7/Epo and REH cells, at days 1, 2, 3, and 4 of culture. Data are expressed as a percentage of β-actin expression relative to day 0.

All results are presented as the mean ± SD of three independent experiments.

### S1 Fig

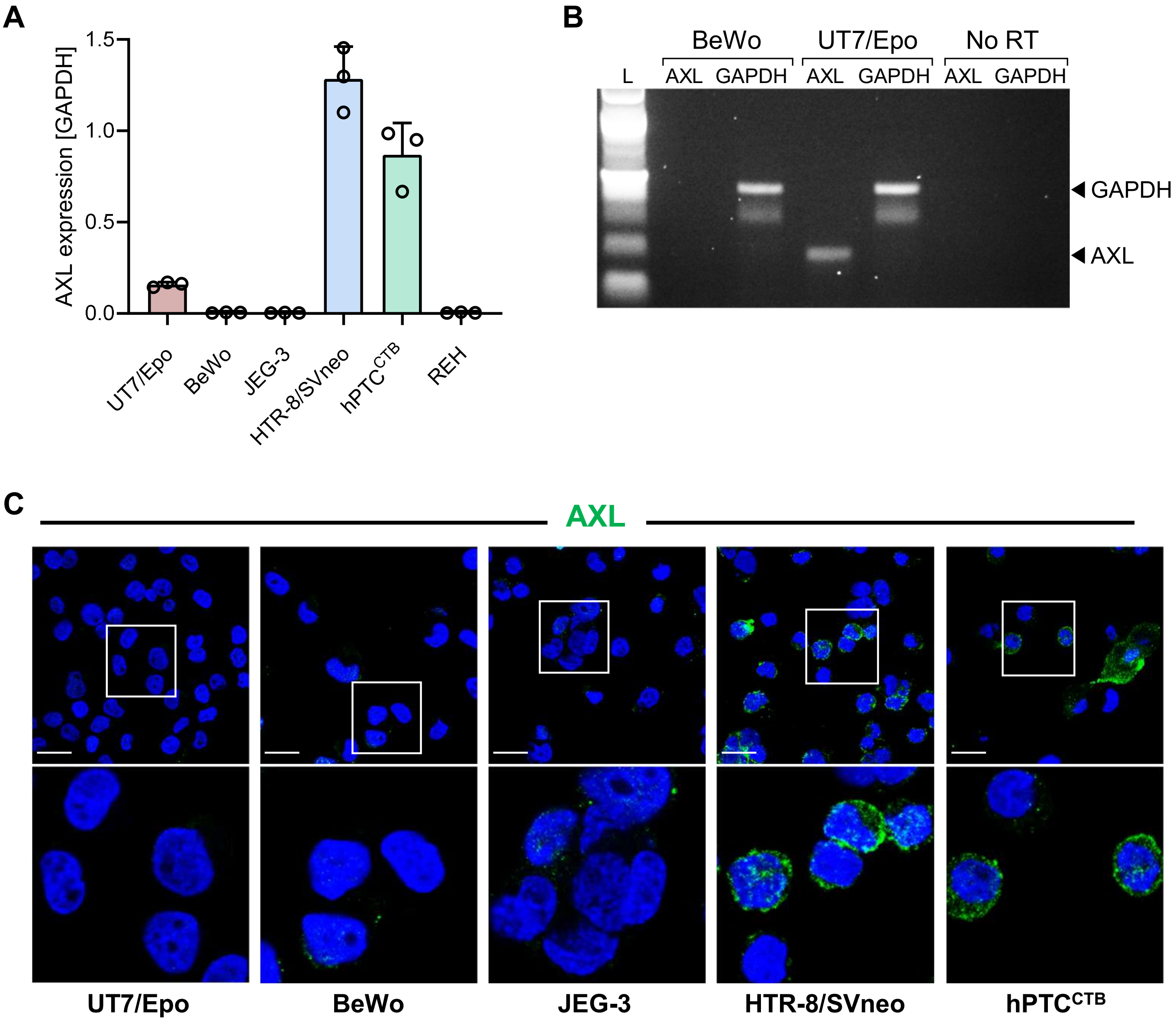
A
AXL expression [GAPDH]
1.5
1.0
0.5
0.0
UT7/Epo
BeWo
JEG-3
HTR-8/SVneo
hPTC^CTB
REH
B
BeWo
UT7/Epo
No RT
L
AXL
GAPDH
AXL
GAPDH
AXL
GAPDH
GAPDH
AXL
C
AXL
UT7/Epo
BeWo
JEG-3
HTR-8/SVneo
hPTC^CTB

### S2 Fig

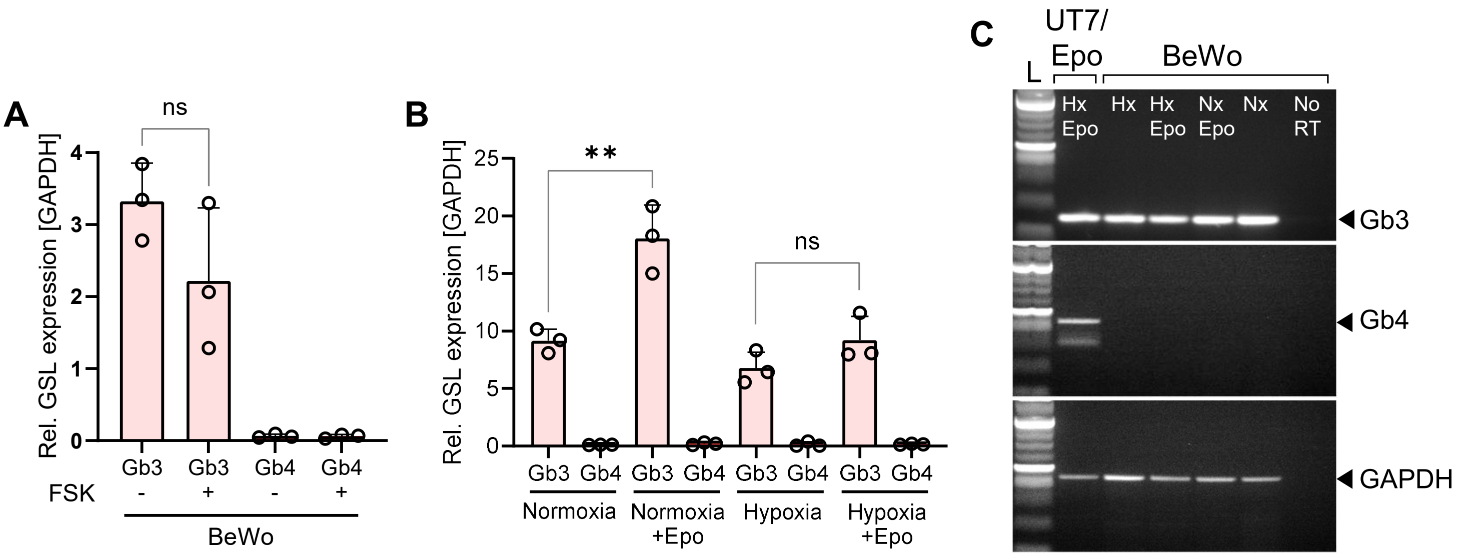
A
ns
Rel. GSL expression [GAPDH]
4
3
2
1
0
Gb3
Gb3
Gb4
Gb4
FSK
-
+
-
+
BeWo
B
**
ns
Rel. GSL expression [GAPDH]
25
20
15
10
5
0
Gb3
Gb4
Gb3
Gb4
Gb3
Gb4
Gb3
Gb4
Normoxia
Normoxia +Epo
Hypoxia
Hypoxia +Epo
C
UT7/ Epo
BeWo
L
Hx Epo
Hx
Hx Epo
Nx Epo
Nx
No RT
Gb3
Gb4
GAPDH

### S3 Fig

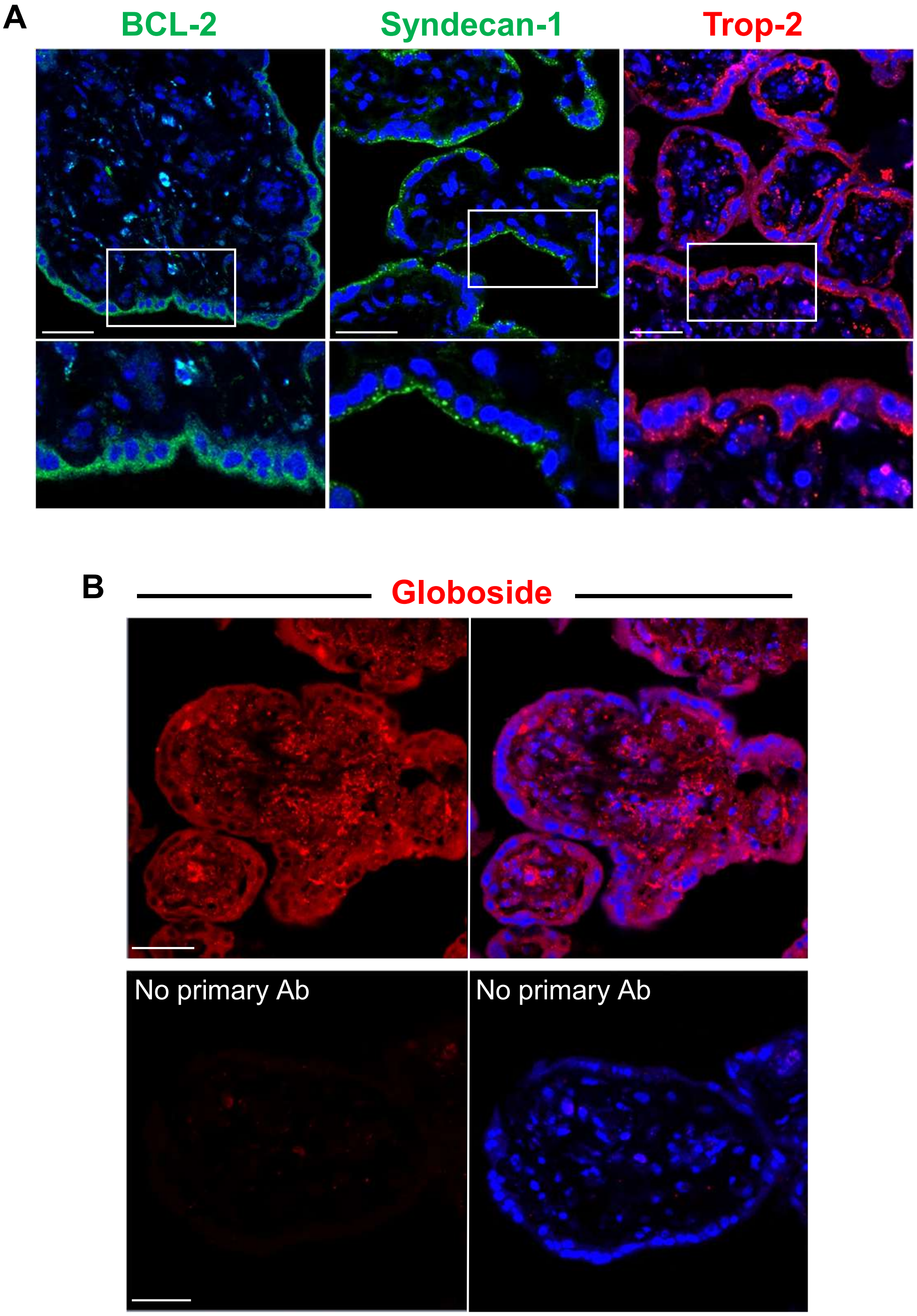
A
BCL-2
Syndecan-1
Trop-2
B
Globoside
No primary Ab
No primary Ab

### S4 Fig

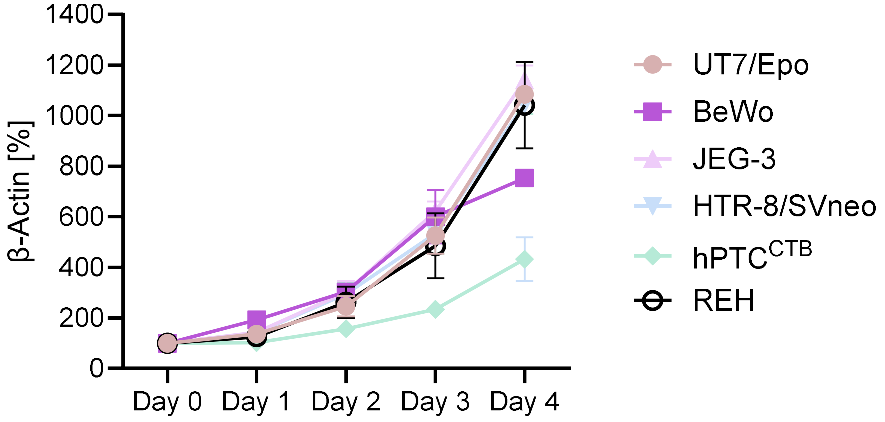

β-Actin [%]
1400
1200
1000
800
600
400
200
0
Day 0
Day 1
Day 2
Day 3
Day 4
UT7/Epo
BeWo
JEG-3
HTR-8/SVneo
hPTC^CTB
REH
